# A Computational Model Captures Orexin-Dependent Modulation of Memory Consolidation

**DOI:** 10.64898/2026.10.02.756298

**Authors:** Polina Shakleina, Adam T. Guy, Enrico M. Trombetta

## Abstract

Hippocampal sharp-wave ripple (SWR) events occur during post-experience rest and sleep and play a significant role in memory consolidation. Recent *ex vivo* studies have found orexin-driven neuromodulatory effects on SWR parameters and proposed that they can influence memory consolidation. Additionally, behavioral studies demonstrate contradictory results regarding orexinergic effects on memory acquisition depending on task structure and intervention time. This inconsistency, along with the diffuse spread of orexin receptors (OXR) in the CA1 and CA3 hippocampus, challenges theoretical interpretation of orexinergic effects on memory. To reconcile this problem, we propose a rate-based model linking orexinergic tone-dependent SWR statistics to memory performance. The model combines an encoding module whose gain scales with the global orexinergic tone with a consolidation module driven by OX2R-specific replay dynamics. The model is based on a plasticity rule that considers SWR events as the unit of induction. Therefore, weight change requires sufficient replay content and depolarizing drive. We observed that, depending on the task structure, encoding-dominant learning benefits discrimination as the orexin drive increases. Consolidation-dominant learning shows a non-monotonic relationship with an internal optimum level of orexinergic tone. This interaction highlights that stronger truncation at higher receptor engagement shortens memory replay and reduces the efficacy of the consolidation process. The model predicts that the optimal regime depends on task complexity reflected in sequence length. Together, these results demonstrate how task demands determine whether increased orexinergic drive enhances or impairs effective consolidation and SWR-dependent replay in the hippocampus.

## Introduction

Orexin (also known as hypocretin) neurons are a small population of excitatory cells in the lateral hypothalamic area (LHA) with a broad impact on brain function. Discovered in 1998 (De Lecea *et al*., 1998; Sakurai *et al*., 1998), orexin peptides exist in two isoforms: orexin A and orexin B. Both peptides bind to two G-protein-coupled receptors, OX1R and OX2R. OX1R is selective for orexin A, while orexin B binds to both receptors with similar affinity (Sakurai *et al*., 1998). Orexin neurons send signals throughout the cortex and subcortical areas (Peyron et al., 1998), which supports their role in controlling arousal, motivation, stress response, and autonomic functions (Sakurai & Mieda, 2011; Barson & Leibowitz, 2017; Soya et al., 2013; Fan *et al*., 2017). Orexinergic stimulation, defined as neuronal modulation conditioned by interaction between orexin peptides and their receptors and subsequent excitation of the afferents, contributes to reward-seeking, emotional response, and alertness rather than directly causing a specific behavioral response (Mahler et al., 2014; Giardino & de Lecea, 2014; Schöne et al., 2014).

In addition to its established roles, orexin signaling is linked to memory processes. In the case of aversive learning, blocking orexin receptors in the locus coeruleus (LC) before training hindered the formation of a conditioned fear response (Sears et al., 2013). Administration of selective OX1R antagonists immediately after training had little or no effect, while pretraining blockade impaired learning, which suggests that orexin A is important for encoding. However, OX2R antagonist injection before training had no effect on the behavior, implying that OX2R signaling is not necessary for the acquisition (Sears *et al*., 2013). Another study showed that orexin–LC signaling participates in the consolidation process (Soya *et al*., 2013). Therefore, we cannot clearly say whether orexins mainly help with learning during the experience, stabilize memories afterward, or do both. In spatial memory tasks, orexins can improve or impair performance depending on the experimental conditions and context. Some studies have shown that orexinergic stimulation can hypercompensate for stress outcomes and disorders including social anxiety and depression, learning difficulties in an epilepsy model and post traumatic Stress Disorder, (Staton *et al*., 2018; Zhao *et al.,* 2014), but does not improve the performance of intact animals (Deadwyler *et al*., 2007; Mavanji *et al*., 2017; Han *et al*., 2020; Soler *et al*., 2021). This rescue effect shows that orexins affect memory in complex, ambiguous ways. There must be a regulatory mechanism that impacts offline rhythmic hippocampal activity, including SWR.

Sharp wave-ripple (SWR) complexes are oscillatory events that emerge during rest and sleep following experience (Girardeau *et al*., 2009). Sharp waves are regulated and synchronized by inputs from the medial entorhinal cortex (mEC). These events propagate the replay of waking neural activity, through which experience-related patterns promote synaptic plasticity and long-term memory storage (Zutshi & Buzsáki, 2023). Experiments using sensory distraction to disrupt post-ripple consolidation showed that intervention immediately after the SWR event during rest periods between trials significantly slows the learning curve in non-spatial tasks, demonstrating that SWRs are critical for immediate processing and early consolidation of experience during active learning. (Nokia *et al*., 2012). Pharmacological disruption of receptor tyrosine-protein kinase ErbB4 signaling altered SWR firing dynamics, resulting in decreased post-training performance on a spatial working memory task, confirming that precise SWR regulation is required for cognitive function. (Robinson *et al*., 2022).

The hippocampus receives sparse orexinergic innervation from the lateral hypothalamus (Riahi *et al*., 2015). The expression of orexin receptors in CA1 and CA3 (Krause *et al*., 2024; Yaeger *et al*., 2024), together with behavioral evidence showing that hippocampal orexin receptor blockade or deficiency impairs spatial and associative memory tasks (Akbari *et al*., 2006; Mavanji *et al*., 2017; Zhao *et al*., 2014), demonstrates how the central orexinergic system actively modulates hippocampus-dependent memory processes. In a recent study, Kostanšek and coworkers showed that *ex vivo* application of OX1R and OX2R antagonists SB-334867 (OX1R antagonist), EMPA (OX2R antagonist), and TCS-1102 (dual orexin receptor antagonist, DORA) alters SWRs. Blockade of either receptor doesn’t have an equal effect on sharp wave incidence, amplitude, and duration. The analysis showed SWRs’ sensitivity to OX2R antagonism: EMPA and DORA increased ripple duration, but the same effect was absent under OX1R blockade. However, endogenous orexin appears to act through OX2R both to promote ripple generation and to restrain ripple duration, which suggests that orexin signaling tunes hippocampal network activity during the offline rest period (Kostanšek *et al*., 2023). The mechanism by which orexin-dependent alterations in SWR incidence, amplitude, and duration shape consolidation remains undefined (Kostanšek et al., 2023). Here we propose a mechanism where orexin tone modulates hippocampal circuits, altering SWR parameters that shape the efficacy and quality of memory replay.

We hypothesized that the inconsistent effect on behavior arises from the influence on multiple components of offline consolidation. SWR-related consolidation could depend on both how synapses are altered during replay and on the replay events’ completeness, defined as how long the replay sequence lasts. Increased neuromodulatory drive should help memory if the replay structure stays intact, but it could harm the consolidation event in case of sudden truncation of the sequence. Based on the interaction between synaptic strength and replay completeness, we predict that the relationship between orexin levels and memory corresponds to an optimal level of orexin modulation, which depends on the task, especially on the length and complexity of the stimuli sequence.

In the current study, we developed a rate-based computational model that links orexin tone, SWR dynamics, and consolidation efficacy. Our model combines an encoding module with gain scaled by global orexinergic tone and a consolidation module in which OX2R-dependent statistics affect SWR-mediated replay and consolidation. We used OX2Rs asymmetry to structure the model. Because ripple duration changed only under OX2R blockade, and because OX2R blockade reduced ripple incidence most strongly, we used the OX2R-selective condition to calibrate the SWR statistics entering consolidation (Kostanšek et al., 2023). Thus, encoding gain was scaled by global orexinergic tone, since pretraining OX1R blockade impairs acquisition whereas pretraining OX2R blockade does not (Sears et al., 2013). Using SWRs extracted from open-source rat local field potential (LFP) recordings (Grosmark & Buzsáki, 2016) together with experimentally measured orexin receptor effects on SWR properties (Kostanšek et al., 2023), we generated synthetic sharp-wave and ripple event libraries and evaluated how orexin-dependent replay dynamics affect consolidation throughout task structures with different sequence demands. We found that in an agent rate-based experiment with increasing orexin drive, encoding-dominant learning showed a monotonic improvement in the discrimination index, whereas consolidation-dominant learning exhibited a non-monotonic curve with an optimal intermediate. The optimal value reflected a trade-off between weak event-level induction at low OX2R, where SPW are smaller, and incomplete replay at high OX2R engagement. The decreased effective OX2R signaling lowers SPW amplitude and therefore the event-level induction gate. The framework further predicted that the optimum shifts systematically with task complexity. Our results provide a mechanistically constrained explanation for how increased orexinergic neuromodulation can facilitate or disrupt memory depending on task structure by simultaneously controlling replay completeness and synaptic potentiation during SWR events.

## Methods

### LFP data, metadata resolution, and state restriction

We used local field potentials (LFPs) from multichannel .lfp/.eeg int16 recordings from a dataset in the CRCNS repository as our biological anchor (Grosmark & Buzsáki, 2016). Session metadata, including channel counts and sampling rates, was extracted from accompanying XML files and confirmed to match the binary file size. We excluded sessions in cases of missing metadata, inconsistent file size, or when sleep-state annotations couldn’t be loaded. One session was excluded after detection due to the small number of detected SWRs. Analysis methods were restricted to the POST epoch and to NREM or Intermediate intervals. Further preprocessing included merging overlapping or adjacent target intervals. Event rates were reported per minute of total included NREM/Intermediate time.

### Channel quality control and laminar-informed channel selection

Because of the inconsistent maximal detection of SPWs and ripples within CA1 laminar depths, we used a representative sample from the first POST NREM/Intermediate period to choose channels independently for SPW detection, ripple detection, and noise turn down. A 300 -second window centered within the timeframe (±150 seconds at the midpoint, trimmed to available data) was extracted for every channel. In order to exclude flat, saturated, or artifact-dominated channels, channels with unrealistic amplitude statistics (standard deviation < 5 or > 5000 ADC units) or high kurtosis (|*kurtosis*| > 10) were eliminated. The top 8 channels were chosen after candidate SPW channels were evaluated according to variance in the SPW band (5–40 Hz). Raw variance in the ripple band (150–250 Hz) was used to rank candidate ripple channels, specifically excluding the SPW channels that had already been chosen. In order to facilitate artifact rejection, four noise reference channels were chosen from the remaining pool by optimizing a noise score (300–500 Hz power relative to ripple power).

### Chunked processing and boundary control

Direct event detection was carried out in 120-s blocks with a 4-s overlap at the native sampling rate (usually 1250 Hz). With the exception of the first and final chunks in an interval, detections were only accepted within the “core” of each chunk (excluding the overlap margins) in order to avoid boundary artifacts from filtering and envelope estimation.

### SPW detection

A multi-channel envelope-based detector was used to find SPWs inside the chosen NREM/Intermediate intervals. SPW-channel data were bandpass filtered (5–40 Hz; 4th order Butterworth; zero-phase SOS filtering) for every block. An aggregate SPW envelope was defined as the mean envelope across SPW channels, and the analytic amplitude envelope was calculated for each channel using the Hilbert transform. Candidate SPW segments were identified where the aggregate envelope exceeded a robust threshold:

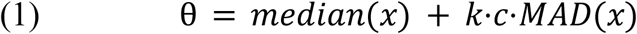

where *k* = 4. 0 and *MAD*(*x*) = *median*(∣*x* − *median*(*x*)∣). In order to guarantee that the *MAD* functions as a reliable estimator of the standard deviation for Gaussian data, the scaling constant *c*≈1. 4826 is defined as 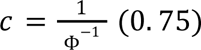, where Φ^−1^ is the inverse cumulative distribution function of the standard normal distribution. Events were kept if they lasted between 30 and 200 ms and if, during the event window, at least two channels surpassed their individual robust envelope limits. Event amplitude was defined as the aggregate envelope at peak, and peak time was defined as the time corresponding to the maximum of the aggregate envelope inside the event. A refractory criterion of 50 ms kept the higher-amplitude event when detections happened in close succession, and events spanning blocks/intervals were combined if they were separated by less than 30 ms.

### Ripple detection with noise-band veto and spectral purity features

A multi-channel envelope detector with an explicit high-frequency noise restriction was used to identify ripples on ripple-selected channels. Hilbert amplitude envelopes were calculated for each channel after ripple-channel signals were bandpass filtered in the ripple band (150–250 Hz; 4th order Butterworth; zero-phase). In order to highlight synchronized high-amplitude ripple activity and lessen susceptibility to single-channel spikes, the aggregate ripple envelope was defined as the 75th percentile of the per-channel envelopes at each time point. Candidate ripple segments that lasted between 20 and 100 ms and had at least three ripple channels that exceeded their respective per-channel envelope thresholds inside the segment were maintained. These segments were found when the aggregate envelope exceeded a robust threshold with *k* = 5. 0.

A noise veto was calculated using noise-reference channels by filtering in a high-frequency noise band (300–500 Hz), calculating Hilbert envelopes, and creating a mean noise envelope across noise channels in order to reject broadband transient distortions. In order to exclude high-frequency transients that emerge concurrently across channels, candidate ripples were eliminated if the highest noise envelope within the ripple window exceeded a robust threshold (*k* = 4. 5).

To impose an oscillatory structure, each event had to have at least four cycles, which were calculated using zero-crossing counts on the mean ripple-band filtered waveform across ripple channels (*cycles* = *zero*_*crossings*/2). Welch’s approach was used to calculate spectral features for each surviving ripple using the raw ripple-channel data in a 160-ms frame centered on the ripple peak (*n_perseg_* = 128, *n_overlap_* = 64). The ratio of integrated ripple-band power (150–250 Hz) to integrated gamma-band power (60–120 Hz) was used to determine spectral purity. To stabilize ratios, gamma power was floored at the fifth percentile of gamma power estimates across sample windows. The PSD maximum within the ripple band was used to define peak ripple frequency. Similar to SPWs, ripple events were combined if they were separated by less than 15 ms, with a 30-ms refractory limit.

### SPW–ripple coupling and SWR-tier definition

Peak times were used to evaluate SPW-ripple coupling. If a ripple peak occurred within a window of −20 ms to +50 ms in relation to each SPW peak, it was deemed linked; if more than one ripple satisfied the window, the closest ripple in time was assigned. Next, two ripple tiers were identified: (1) All ripples: events that met spectral limitations (spectral purity ≥ 1.5; peak frequency 140–220 Hz); and (2) SWR-tier ripples: coupled ripples that also met more stringent requirements (spectral purity ≥ 2.0; peak frequency 150–200 Hz). The percentage of SPWs with a designated ripple in the coupling window was used to report coupling rates.

### Detector validation and quality control

Detector performance was validated by visual inspection of randomly sampled SWR-tier events showing wideband LFP, SPW-band activity, ripple-band activity, and local spectrograms aligned to ripple peaks; and pooled distribution checks of SWR-tier duration, peak frequency, spectral purity, and coupling lag distributions across sessions. Session-level QC metrics included ripple-channel coverage, SWR-tier median spectral purity, and coupling-lag dispersion. For downstream modeling, high-confidence subsets were defined as SPWs with 50–120 ms duration and amplitude in the upper 50th percentile (and ≥2 channels), and SWR-tier ripples with 20–80 ms duration, power in the upper 50th percentile, and ≥2 channels (additionally maintaining the SWR-tier spectral purity and peak-frequency constraints).

### Synthetic SPW–ripple generator and orexin modulation

Synthetic SPW and ripple event libraries were generated from empirical distributions estimated from the validated detector output. For each session, SPW and SWR-tier ripple features (durations, amplitudes/powers, peak frequencies, and coupling lags) were extracted from POST NREM/Intermediate intervals. Inter-event intervals (IEIs) were computed within continuous NREM/Intermediate intervals to avoid cross-gap artifacts.

Empirical marginal distributions were fit using parametric right-skewed distributions. SPW duration, SPW IEI, ripple duration, and ripple IEI were modeled with Gamma distributions. SPW amplitude was modeled as lognormal, supported both by the established lognormal organization of hippocampal SPW-R magnitudes (Mizuseki & Buzsáki, 2013; Buzsáki & Mizuseki, 2014) and by within-dataset comparison (Δ*AIC* = 295. 8 favoring lognormal over Gamma). For ripple power, no comparable literature precedent exists, so the family was selected on data alone using AIC; in the present dataset, Gamma provided the better fit (Δ*AIC* = 17. 8 vs. lognormal). Both Gamma and lognormal fits, AIC values, and Kolmogorov–Smirnov statistics are reported in Supplementary Fig. 4C. Coupling lags were sampled from the empirical lag distribution, with Gaussian noise scaled by a model-specific precision parameter and values constrained to the coupling window.

For each 3-s trial, SPW peak times were drawn from a homogeneous Poisson process with rate:

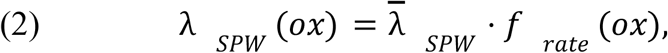

and each SPW of amplitude *a* was assigned a ripple with probability

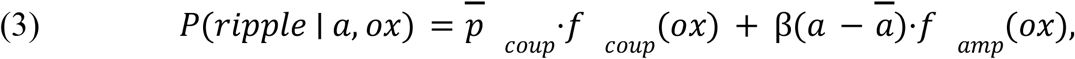

Where *λ̅_SPW_*, *p̅_coup_*, β, *̅a* are empirical baselines (SPW rate, coupling probability, amplitude–coupling slope, and median SPW amplitude), and *f_rate_*, *fcoup*, *f_amp_* are orexin-model-specific scaling factors. For coupled SPWs, the number of ripples was sampled from the empirical ripples-per-SPW distribution reweighted by a model-dependent multi-ripple tendency. Each ripple’s frequency *f_r_* and cycle count *n* _c_≥3, below the detector’s four-cycle criterion, were drawn from empirical distributions, with duration *d* = *n_c_*/ *f_r_* capped at 250ms. Ripple power was sampled independently. Uncoupled background ripples were added as a Poisson process at a fixed fraction of α(*λ̅_ripple_*_, *all*_− *λ̅_SWR_*), with α = 0. 5 as a fixed fraction.

Orexin manipulations were implemented as parametric modulation models controlling SPW rate/amplitude/duration, ripple duration, and coupling probability as functions of a scalar orexin level. Paper-aligned models used a ripple frequency band of 100–175 Hz, while exploratory models used 150–250 Hz, matching the frequency bands used for downstream comparisons. Synthetic libraries were generated on a dense 15-level orexin grid spanning 0.3–1.8, with 200 trials per model × context × orexin-level condition. To model the antagonists’ effects, the multipliers were taken from the changes reported by Kostanšek et al. (2023). Three antagonists (SB-334867 (OX1R), EMPA (OX2R), and TCS-1102 (DORA)) were implemented as separate modulation models, each specifying multipliers on sharp wave rate, amplitude, and duration, ripple duration and power, and coupling probability, defined so that ox = 0.3 reproduces the reported drug condition and ox = 1.0 reproduces baseline, with linear interpolation between (Table 1). The consolidation analysis used the EMPA-derived libraries, while the SB-334867 and TCS-1102 models were not used in the agent simulations and remained in the report as a calibration control. Orexin’s level here is a dimensionless index of effective OX2R signaling normalized to the intact condition. The axis is anchored at two points: ox = 1.0 reproduces the baseline SWR statistics of the source recordings, and ox = 0.3 reproduces the statistics measured after bath application of EMPA (Kostanšek et al., 2023); intermediate levels interpolate linearly between them. Because the calibration derives from antagonism, modulation saturates at ox = 1.0, and library statistics are identical for all supra-baseline levels; effects above baseline therefore arise from the consolidation-side termination mechanism and are presented as model predictions.

### Sequence-based memory task construction

We modeled episodic memory as a discrimination between two traces sharing a common beginning and diverging partway through. Each trace was a single *d*-dimensional vector (*d* = 64) in which dimension index stands for position along the remembered sequence: the first *n_s_*_ℎ*ared*_ = ⌊*d* · *pdiv*⌋ dimensions carry the shared prefix, the remainder carry the trace-specific continuation. At the divergence fraction used throughout, 38 dimensions are shared, and 26 are unique. Components were drawn independently from a standard normal distribution, and each assembled vector was normalized to unit length. Trace A was assigned outcome 1 and trace B outcome 0, which confines discriminative information to the post-divergence block.

Sequence structure initiates in the model at the replay stage rather than at encoding. Each replay event moves along the memory vector that begins at dimension 0 and stops wherever the event terminates, so a truncated event exposes the first part of the trace and hides the rest. Because the shared block occupies the first 38 dimensions and the trace-unique block the last 26, the point at which discriminative information first becomes available is fixed: it is the step at which the walk finishes the shared block, ⌊*seq*_*len* · *n_s_*_ℎ*ared*_/ *d*⌋ = 9 of a nominal 16.

Events terminating at or before step 9 reveal only shared dimensions and therefore carry no information distinguishing the two traces, however strong the plasticity acting on them. Beyond step 9, the unique block is revealed in proportion to the remaining steps, reaching full exposure at step 16. Replay quality *q*, the fraction of unique dimensions revealed, is the model’s measure of this and enters the plasticity gate directly (see *Consolidation model*); *q* is zero up to and including step 9 and rises to 1 across steps 10 to 16 (Supplementary Fig. 1).

In order to evaluate the agent’s ability to discriminate between two traces A and B, they are both evaluated 20 times; at each iteration, with added Gaussian noise (SD = 0.02), each noisy readout was clipped to [0, 1], and DI was the difference between the two means.

### Encoding model

Each replicate began with a new agent, so no weights carried over between conditions and there was no pre-training stage. The agent uses a single linear weight vector 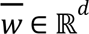 with logistic output, 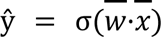 with x being the full unit-normalized d-dimensionalized trace, the pre-activation clipped to ±30 for numerical stability. Weights were initialized from a zero-mean Gaussian with variance 2/*d* (He et al., 2015). The decision boundary passes through the origin; therefore, no bias term was included. Because traces are unit-normalized and *w̅* is initialized independently, both traces are read out near ŷ = 0.5 before learning, and discrimination is initially zero. During encoding, *x̅* is the full trace; during consolidation, it is the same trace after truncation and additive noise (see *Consolidation model*). Thus, partially replayed events both expose fewer dimensions and lower readout drive. During EC, weights change according to a delta rule,

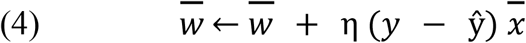

which used only the presented activity pattern and a scalar error at the output unit. The readout had no hidden layer, so no gradient was propagated through it.

Awake encoding was repeated presentation of each trace with its outcome, at learning rate.

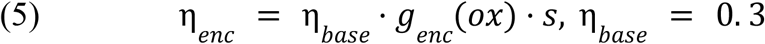

where *s* is a per-trace encoding strength, and *g*_enc is a saturating function of global orexinergic tone, standing in for the OX1R-like pathway:

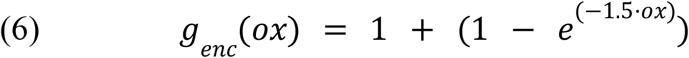

The gain rose monotonically from 1 with no orexin drive toward an asymptote of 2, spanning 1.36 to 1.93 across the tested range of 0.3 to 1.8. The target trace was presented 10 times at *s* = 0. 5, the lure 3 times at *s* = 0. 15. Discrimination after encoding was therefore positive but short of ceiling, which left room for consolidation to move it in either direction.

The two pathways read different inputs. Encoding saw only the scalar orexin level and never touched the synthetic SWR library; consolidation read the library and nothing else. In encoding-dominant runs, the tone term tracked the swept level. In consolidation-dominant runs, it was held at zero for every level, fixing (6) at its floor of 1, so that all variation arose in consolidation. We used a common random numbers method, so the encoding arm contains no stochastic component varying with OX2R effective activation level. Therefore, the rank order of DI within a replicate is fixed by (6). Neither pathway was fitted to produce dissociation; however, they are asymmetrical in their architecture by design. Because (6) appears as a saturating monotone function, the encoding can only produce a monotone DI response, which is set by the gain function.

The delta rule updates weights toward an outcome target, an abstraction of an outcome-associated readout receiving feedback about the remembered episode; the results below do not depend on the learning rule itself being biologically exact.

### Consolidation model

Since the varying of the effective OX2R activation affects the length of the replay period, our model required an induction to depend on replay progression instead of event count alone.

Consolidation is gated at the level of individual SWR events by a three-factor rule in which the content, the timing, and the drive of an event each had to be adequate for any weight change to occur. We model plasticity through the SPWs’ amplitude, which influences consolidation by increasing the probability of induction while simultaneously accelerating the accumulation of inhibitory signals that terminate the replay process. The magnitude of induction depends on the product of these two parameters, scaled by the degree of coupling between the ripple and the sharp wave, and becomes zero if this product falls below a specified threshold. Rising OX2R moves both effects, scaling the inhibitory accumulation rate through k(ox) while shifting the amplitude and event-rate statistics that set the drive.

The replay quality *q,* defined as the fraction of post-divergence steps completed before termination, is scaled to [0, 1] and raised to a truncation exponent γ (γ = 1 in the core model). Events terminating at or before the divergence step had *q* = 0. The replay quality sets the fraction of memory-unique dimensions revealed to the readout. Therefore, events too short to pass the divergence point carried no discriminative content regardless of their other properties.

The coincidence and external SPW drive factors formed the LTP gate *g*. We modeled coincidence as a Gaussian in SPW–ripple coupling lag, ^(−*lag*²/2*w*²)^ with *w* = 10 ms, imposing maximal induction to tightly coupled events. Sharp waves in a block without a coupled ripple were assigned zero lag. The kernel has the functional form of an STDP window, but its argument is the lag between the sharp wave and the ripple it carries, not the interval between pre- and postsynaptic spikes. Plasticity induction during ripples has been demonstrated directly (Sadowski et al., 2016); the kernel expresses the assumption that induction is most effective when the two components of the event are tightly aligned. The drive factor was the logistic of SPW amplitude *z*-scored against the baseline library distribution, treating amplitude as a proxy for the depolarizing drive available during the event. Induction was then a thresholded, supralinear function of the combined signal *s* = *q*·*g*:

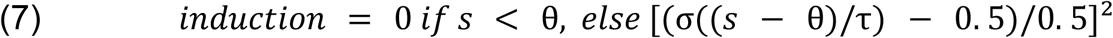

with θ = 0.20 and τ = 0.05. Amplitude normalization is calibrated from the baseline library. The induction threshold and slope, the consolidation rate ratio, and the truncation exponent each were fixed at a neutral value and varied singly in the robustness battery.

The threshold plays the role of a modification threshold in the sense of BCM theory (Bienenstock et al., 1982), in which the sign of plasticity is set by whether postsynaptic drive falls above or below a criterion, and that criterion itself moves with the neuron’s recent activity. In our framework, the rule was rectified rather than sign-changing: weights have already changed sign through the error term of the delta rule, so a depression branch below threshold would introduce a redundant second source of sign. Since the model contained no homeostatic regulation of the induction criterion, θ was held fixed rather than sliding with recent activity following calcium threshold, in which a fixed induction criterion is sufficient to produce bidirectional plasticity without a moving threshold (Shouval et al., 2002). The requirement that events exceed a minimum drive before any potentiation occurred corresponded to the cooperativity of LTP induction (McNaughton et al., 1978), and the supralinear growth above threshold to the cooperative calcium dependence of induction (Graupner & Brunel, 2012). Squaring the re-centered logistic made the onset C1-smooth, so that the transition at θ introduced no discontinuity in slope. The per-event learning rate was η₀ · *s*_*consol* · *induction*, with η₀ = 0. 3 and *s*_*consol* = 0. 20, bounding the applied step at 0.06. The decomposition into η₀ and s_consol keeps η₀ shared with the encoding module, so that the free consolidation parameter is a ratio: a maximally induced replay event moves the readout one fifth of the base learning-rate step. Because memory vectors are unit-norm and the readout is trained by exact gradient descent on log-loss, the largest eigenvalue of the loss Hessian is bounded by *p*(1 − *p*)‖*x*‖² ≤ 1/4 and updates are stable for η < 8. The applied maximum is approximately 130-fold below this bound, so the concave shape reported below cannot arise from step-size instability at high OX2R.

Thus, the model isolated the interaction between truncation and event-level induction. Every planned event was eligible for replay, and the planned event count followed SPW incidence carried by the library. Replay fidelity was held at its baseline value, fixing the noise multiplier at 1. A supplementary ablation model enabled OX2R-dependent coupling probability and fidelity-scaled noise, both interpolated from the library, and switched four mechanisms independently under common random numbers: truncation, coupling probability, fidelity, and the LTP gate. When the LTP gate was disabled, g was replaced by a calibrated fixed value. Therefore, a fully completed replay clears the induction threshold, since any constant below θ would leave induction identically zero and make the frozen arm consolidate nothing at every OX2R level. The difference between this arm and the intact model gives the LTP contribution shown in Fig. 4C.

### Replay sequences

Consolidation occurs during offline replays of the stored traces, driven by the synthetic SWR library and is constrained by a fixed plasticity budget expressed in ripple cycles. Weight changes followed the same delta rule as encoding. The two modules differed in what set the learning rate and in how much of each trace was available to be learned.

For each replicate, a replay plan was generated in advance, so that common random numbers aligned across orexin levels. A consolidation window of 5 min was divided into 100 blocks of 3 s, matching the trial length of the generator, and one library trial was drawn per block with replacement. Every planned event inherited its properties from the library trial at the orexin level being simulated: the number of SPW events in the block (and the corresponding ripple cycle count); an SPW–ripple coupling lag; and an SPW amplitude, drawn in shuffled order from that trial’s own values and recycled if exhausted. Cycle counts were floored at the generator’s minimum. Taking the event count from the orexin-specific trial rather than from baseline carried the measured reduction in SPW incidence under OX2R blockade (Kostanšek et al., 2023) into the model. At the lowest level simulated, a plan held roughly half as many events as at baseline.

Replay length was set by a race between excitation and inhibition within each event. Inhibition accumulated in proportion to the normalized SPW amplitude â at a rate *k*(*ox*) = *k_slope_* · *ox*, with *k_slope_* being set to 0.06 as our operating point except where otherwise stated, while excitation decayed by 6% per step from an initial value of 1 perturbed by Gaussian noise (SD 0.05):

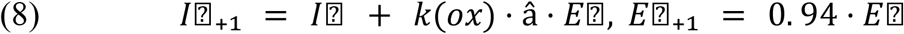

*k_slope_* is a free structural parameter: Kostanšek et al. (2023) constrain LFP signatures rather than replay content, so nothing in the calibration data bounds it.

The event terminated at the first step where *I* − *E* exceeded 0.30, and otherwise ran to the full 16 steps. Larger amplitudes and higher orexin levels therefore produced earlier termination. Cycle counts did not set replay length; they were the currency of the consolidation “budget”. The budget was a fixed total number of ripple cycles available across the whole consolidation window, computed once from the baseline library and held identical at every OX2R level. The truncated trace was then exposed to the readout, with shared dimensions revealed in proportion to progress toward the divergence step and post-divergence dimensions only beyond it, and with independent Gaussian noise added to the visible entries (SD 0.03 × *u*, *u* ∼ U[0.02, 0.06] per event).

Each executed event cost its cycle count scaled by completeness, so truncated replays drew less from the budget, and an event was skipped when its cost would exceed the remaining budget. That ceiling was the mean total cycles per baseline trial multiplied by the number of blocks and divided by a target utilization of 0.90. The trace that was replayed on a given event was drawn in proportion to encoding strength, giving the target about 77% of events.

### Simulation design and statistical reporting

Simulations were performed as Monte Carlo sweeps across effective OX2R activation levels using multiple independent replicates (50 for the main, frozen-gate and truncation-off sweeps; 25 for the dual-pathway, divergence, complexity, robustness and k_slope sweeps; 30 for the encoding-sensitivity and consistency analyses). All sweeps used common random numbers: within each replicate, the memory pair, the sampled sequence of library trials, and all per-event random draws were generated from a fixed seed and reused across every OX2R level, so that differences between levels cannot be attributed to replicate sampling variability. Two grids were used, a dense nine-level grid spanning OX2R = 0. 3 – 1. 8 for the main curves and the complexity analysis, and a coarse five-level grid (0.3, 0.7, 1.0, 1.3, 1.8) for the robustness sweeps, where the number of conditions made the dense grid impractical.

To disentangle encoding vs consolidation-driven effects, we ran two regimes: an encoding-dominant regime in which orexin tone varied only during encoding, and a consolidation-dominant regime in which encoding tone was fixed at the gain floor and OX2R varied only during consolidation. A third, dual-pathway sweep varied both pathways together, encoding alone with consolidation held at OX2R = 1. 0, and consolidation alone with encoding held fixed, so that the contribution of each pathway to the combined curve could be read directly.

Encoding strength was fixed a priori at 10 presentations, which leaves DI without replay short of ceiling. Because this choice affects the outcome measure, the sweep was repeated at 3, 5, 8, 10, 12, 15, and 20 presentations; q₂ stayed negative, and the vertex stayed between 0.971 and 0.974 across all seven, which keeps the trace below saturation and leaves consolidation able to move DI in either direction. The encoding-dominant sweep used a fixed 12 presentations.

Task complexity was varied in two ways. Sequence length was set from 8 to 28 steps with the divergence fraction held at 0.6 (Fig. 5A), and the divergence step was varied from 3 to 12 at a fixed sequence length of 16 (Fig. 5B).

Mechanistic contributions were isolated by orthogonal ablations using common random numbers. In the frozen-LTP arm, the event-level plasticity gate was replaced by a constant floor while truncation remained intact; this arm was run on the dense grid against the intact model, and the difference between them gives the LTP contribution shown in Fig. 4C. Two further ablation sets, a 2×2 crossing of truncation and plasticity gating and a comparison of the coupling and fidelity modulators, were run with baseline replay plans rather than OX2R-specific ones, and served as internal consistency checks (not shown).

Parameter dependence was assessed by repeating the consolidation sweep on the coarse grid while varying single parameters: readout dimension (32–128), induction threshold θ (0. 10–0. 30) and slope τ (0. 02–0. 20), consolidation learning-rate scale (0.10–0.40), the three E/I termination parameters (κ 0.00–0.12, threshold 0.15–0.40, decay 0.03–0.10), and the truncation exponent γ (0. 5–2. 0). A further control replaced strength-weighted memory selection during replay with uniform sampling. Unlike the library parameters (Table 1), the *E*/*I* termination parameters are not constrained by Kostanšek et al. (2023). Their ranges were chosen to span distinct regimes rather than a calibrated interval. This means that for κ, the *k_slope_* range includes values at which truncation is negligible and values at which most events terminate before the divergence point, so that the contribution of truncation to the descending limb can be read from the sweep itself.

The DI–*OX*2*R* interaction in consolidation sweeps was summarized as

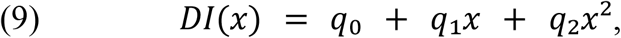

where x denotes effective OX2R activation, and q₂ is the quadratic coefficient. The sampling distribution of q₂ was obtained by resampling replicates 10,000 times. We report q₂ with its 95% percentile interval and the bootstrap sign probability *P*(*q*₂ ≥ 0). Monotonicity was assessed for monotonicity sweeps using Spearman rank correlation between OX2R and DI.

Peak location was estimated as the stationary point of the fitted quadratic, 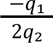, the standard turning-point estimator for a second-order response surface (Box & Wilson, 1951). A continuous estimate took the vertex of the fitted quadratic, identifying intervals by percentile bootstrap over 5,000 resamples. The vertex was clipped to the sampled range and assigned to the nearer endpoint, reporting 95% percentile intervals. A discrete estimate, P(interior), recorded the proportion of draws in which the largest mean DI fell at a level other than the two endpoints. Draws with vertices outside the sampled range accumulate at the boundaries, so intervals whose limit coincides with an endpoint should be read as censored rather than as located.

Because concavity alone is also compatible with a monotonic saturating curve, non-monotonicity was not inferred from the sign of q₂ but from the fitted vertex falling strictly inside the sampled range, together with P(interior). For encoding sweeps, monotonicity was assessed using Spearman rank correlation between OX2R and DI.

The dependence of peak location on task complexity was tested by fitting the model.

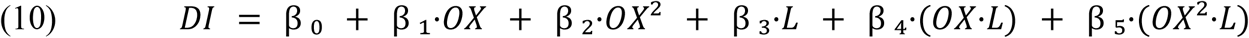

with sequence length (L) mean-centered. For each of the 5,000 bootstrap samples, the estimated peak position shift was calculated based on replicate indices. Within each sample, the peak position was regressed against sequence length. A similar interaction model was applied to the analysis of the divergence point shift, with the depth axis expressed in terms of replay steps.

We tested the encoding/consolidation dissociation as a single contrast. DI was modeled jointly across both arms of the dual-pathway sweep as

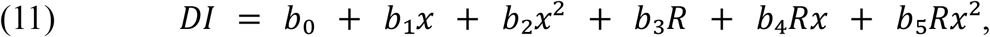

where x is the mean-centered OX2R level, and R indicates the consolidation arm. The coefficient b₅ is the difference in quadratic coefficient between arms and is invariant to translation of x. Because both arms were generated from the same replicate seeds, the assumption used a paired bootstrap in which each resampled replicate index was applied to both arms. The two arms differ in level, because the encoding arm runs consolidation at OX2R=1.0 while the consolidation arm holds encoding at its gain floor. That offset loads on b₃ rather than on b₅, and the contrast was additionally verified with DI standardized within arm, which removes differences in level or scale between them. Point estimates are the fit to all replicates; intervals are 95% percentile intervals over 10,000 bootstrap resamples.

Additional mechanistic diagnostics included P(pre-div) and total replay steps as measures of truncation exposure and functional replay capacity. Other parameters included orthogonal ablations of the plasticity gate, replay truncation, and SPW–ripple coupling. To isolate their separate contributions, we used content fidelity. All reported means are accompanied by standard errors across replicates unless otherwise specified, and all random processes were seeded deterministically per replicate, with the same seed used across OX2R levels so that comparisons between levels are paired.

## Results

### Empirical SPW and ripple extraction and detector validation

Across the three training sessions, the detector identified 6, 619 SPWs and 5, 382 ripples that passed the general ripple criteria. Of these ripples, 2, 054 were temporally coupled to an SPW and satisfied the strict SWR-tier spectral criteria. SPW rates were 26. 1, 15. 9, and 18. 8 events min⁻¹. Corresponding SWR-tier rates were 8. 0, 4. 6, and 6. 3 events min⁻¹. Overall, 31. 0% of detected SPWs were associated with an SWR-tier ripple. A more stringent high-confidence subset retained 2, 197 SPWs (33. 2% of all SPWs) and 982 SWR-tier events (47.8% of the SWR-tier population).

The detected events showed consistent time-based and frequency-based organization. Median SPW-to-ripple peak lags were 0. 0 ms, − 4. 8 ms, and − 4. 0 ms. The pooled lag distribution was centered close to the SPW peak, with a median of −3.2 ms, rather than accumulating at the limits of the imposed coupling window. Across all 2, 054 SWR-tier events, ripple peak frequency averaged 173 ± 13 Hz and remained concentrated within the predefined 150–200 Hz SWR-tier range.

As an extra safety check, thirty SWR-tier events from each session were randomly sampled, and generally they showed both a low-frequency sharp-wave deflection accompanied by a transient ripple-band oscillation, and localized high-frequency spectrogram power around the detected event. The representative event shown in Fig. 1B had a duration of 46 ms, a peak frequency of 176 Hz, and a spectral-purity ratio of 3. 2. The event yields, channel-selection controls, temporal-coupling distributions, spectral measurements, and visual inspection supported using these extracted events as the empirical baseline for the synthetic SWR generator.

**Figure 1.**
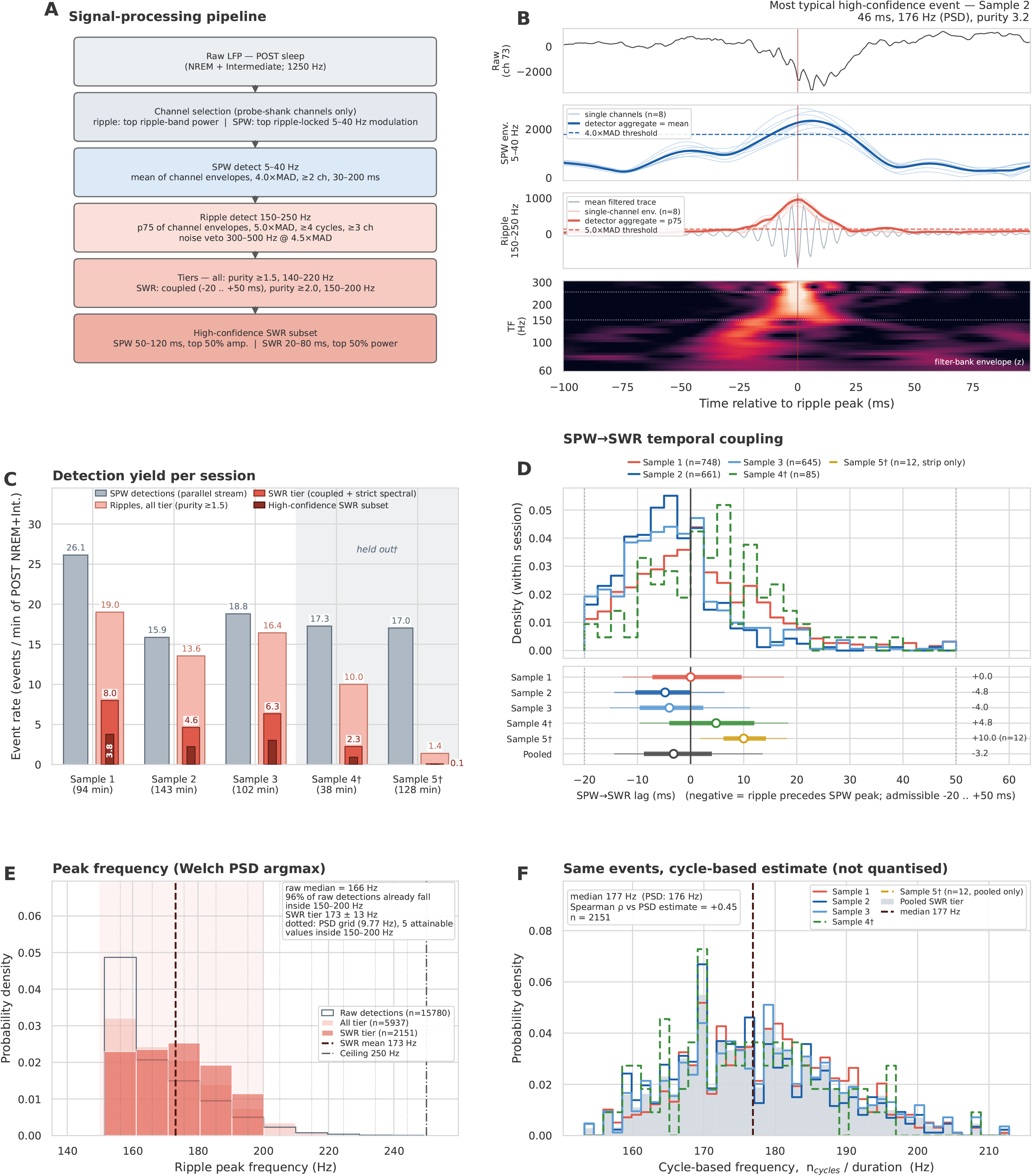
Detection and physiological validation of sharp-wave ripples in open-source hippocampal LFP. **(A)** Detection pipeline, applied to NREM and Intermediate epochs of POST sleep from five sessions (Samples 1–5; three training, one held out, one excluded) at 1250 Hz. Sharp waves: mean 5–40 Hz envelope across channels, 4.0 × MAD, ≥2 channels, 30–200 ms. Ripples: 75th-percentile 150–250 Hz envelope, 5.0 × MAD, ≥4 cycles, ≥3 channels, with events vetoed at 300–500 Hz power above 4.5 × MAD. Nested tiers follow: *all* (purity ≥1.5, peak 140–220 Hz) and *SWR* (coupled within −20 to +50 ms, purity ≥2.0, peak 150–200 Hz). Thresholds are read from the stored detector configuration at figure time. **(B)** Representative event from Sample 2, selected as the high-confidence event closest to the pooled medians of duration, peak frequency, and purity (46 ms, 176 Hz, purity 3.2). Thin traces, single-channel Hilbert envelopes (n = 8 per band); bold traces, the across-channel aggregates of the detector thresholds (mean for sharp waves, 75th percentile for ripples); dashed lines, the corresponding 4.0 × and 5.0 × MAD thresholds recomputed from a 60 s local baseline. Bottom, constant-Q filter-bank envelope, z-scored within the band. **(C)** Yield per minute of POST NREM + Intermediate (93.5, 142.6 and 101.7 min; 37.7 and 127.8 min). Training sessions: sharp waves n = 6,619; ripples all tier n = 5,382, SWR tier n = 2,054, high-confidence n = 982. All five sessions: 9,447, 5,937, 2,151 and 1,021. **(D)** SPW→SWR lag, normalized within session; negative values indicate ripple before sharp-wave peak. Dotted lines: the admissible window, which truncates the distribution by construction. Below, median (open circle), IQR (thick bar), and 10th–90th percentile (thin line) on the same axis. Medians: +0.0 ms [IQR −7.2, +9.2], −4.8 ms [−10.4, +0.0], and −4.0 ms [−9.6, +1.6] for Samples 1–3 (n = 748, 661, and 645); pooled −3.2 ms [−9.6, +3.2]. Sample 4† +4.8 ms (n = 85); Sample 5† +10.0 ms (n = 12, shown as a strip only). Across sessions, 31.0% of sharp waves were followed by a qualifying ripple (2,054/6,619). Lags are resolved to the 0.8 ms LFP sample period. **(E)** Peak frequency. Open steps, raw detections (n = 15,780); filled, all tier (n = 5,937) and SWR tier (n = 2,151), all five sessions. The estimator is quantized to 9.77 Hz (dotted grid), leaving five attainable values in 150–200 Hz, so the discreteness is a property of the measurement. Edge density reflects the 150 Hz detection floor. Raw median 166 Hz, with 96.4% of raw detections already within 150–200 Hz; SWR tier 173 ± 13 Hz (mean ± s.d.). **(F)** The same events scored by cycle count divided by duration, which is independent of the spectral grid. The median is 177 Hz against a PSD median of 176 Hz, confirming that the banding in **E** is imposed by the estimator; single-event agreement is moderate (Spearman ρ = +0.45, n = 2,151), since the cycle measure averages over the event while the PSD argmax tracks its dominant component.

### Synthetic SPW–ripple generator preserves empirical event statistics and pharmacological constraints

The synthetic generator reproduced the pooled empirical SPW and SWR activity that corresponded to the null-model baseline (Fig. 2A). Across 250 min of simulated NREM, the model generated 19. 68 SPWs min⁻¹, 6.27 SWRs min⁻¹, and an SPW-to-SWR coupling probability of 0.319, compared with training-session time-weighted estimates of 19. 59 SPWs min⁻¹, 6. 08 SWRs min⁻¹, and 0. 310, respectively. These corresponded to deviations of 0. 4% for SPW rate, 3. 2% for SWR rate, and 2. 7% for coupling probability. Because SPW rate and baseline coupling were specified during model construction, their agreement confirmed correct calibration.

**Figure 2.**
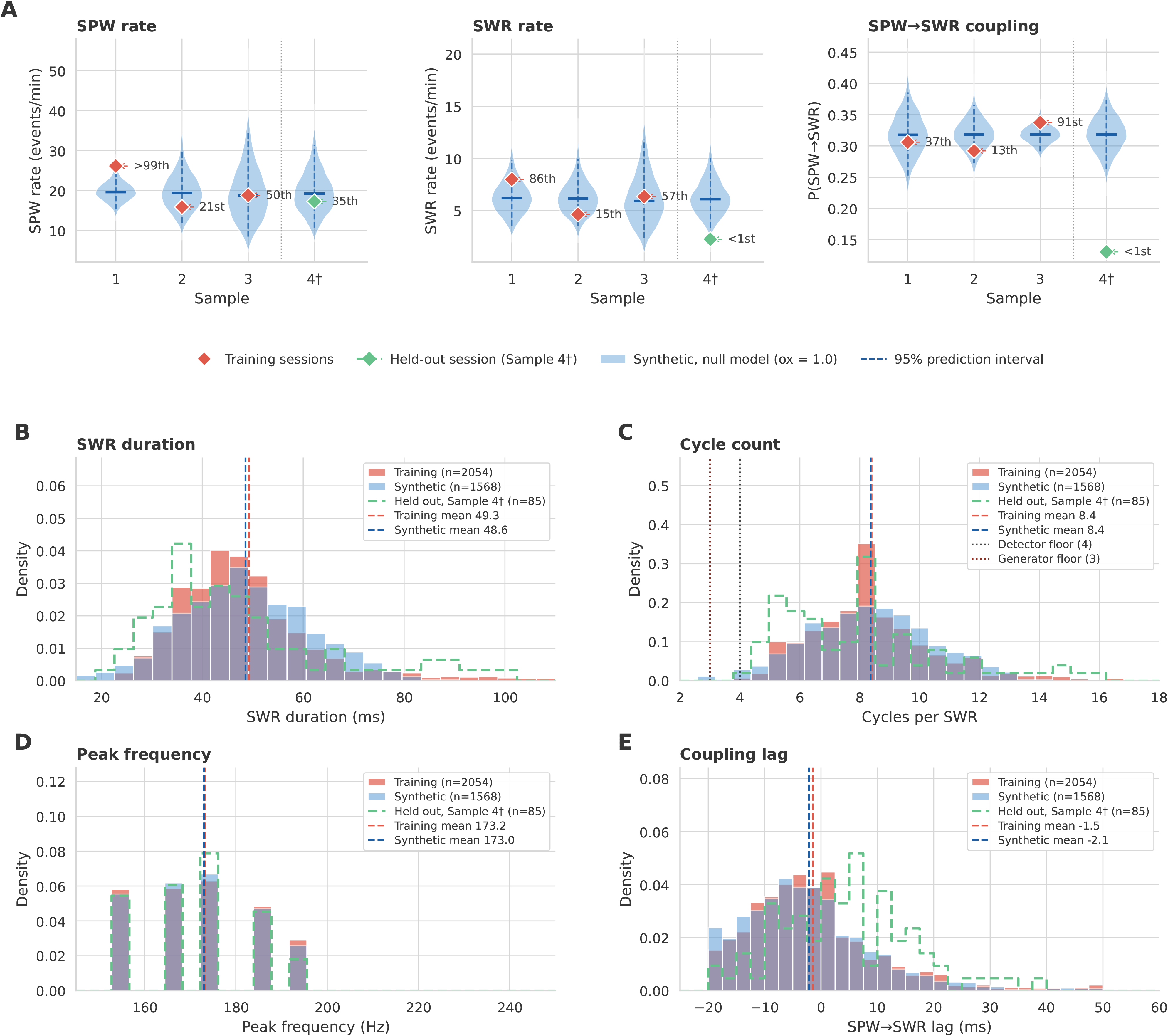
Validation of the synthetic SWR generator. The synthetic library is the null model at ox = 1.0 (5,000 three-second trials, 1,568 SWR-tier events, seed independent of the agent libraries), built from Samples 1–3 only. Sample 4 (†) was held out; Sample 5 is not shown, its ripple detection having failed quality control (Methods). **(A)** Event rates and SPW→SWR coupling per session (diamonds) against prediction intervals for a synthetic session of the same duration (violins; dashed line, 95% interval), with overdispersion set to the larger of the null library’s and that of the other training sessions. Percentiles show each session’s position within its matched distribution. **(B–E)** Pooled event distributions. Training and synthetic are filled, the held-out session is a dashed outline, and dashed vertical lines show the mean. KS statistics are computed on samples matched for n and are reported against a resolution floor, the KS expected between two halves of the same sample at that size. Peak frequency is quantized to the 9.77 Hz Welch grid (Fig. 1E), which produces the discrete bars in D. Panel C carries two floors: the detector requires four cycles, the generator three.

The null model also preserved the principal event-level distributions used to construct replay events (Fig. 2B-E). Mean ripple frequency was nearly identical between empirical and synthetic events (173. 19 vs 172. 98 Hz), as were mean cycle counts (8. 39 vs 8. 36 cycles). The synthetic coupling-lag distribution remained centered near the SPW peak, although its mean was shifted slightly earlier than the empirical distribution (− 2. 11 vs − 1. 45 ms). The mean SPW amplitude was also similar between the datasets (2, 699. 9 vs 2, 697. 3 detector units). The synthetic ripple-duration distribution was close to the empirical distribution, with means of 48. 58 and 49. 25 ms, respectively, and an Earth mover’s distance of 3. 05 ms.

Distribution comparison supported differences between parametric forms for SPW amplitude and ripple power. Compared to a Gamma model, which performed better for ripple power by 17. 8 *AIC* units, a lognormal model provided lower AIC for the amplitude ( Δ*AIC* = 295. 8) (Supplementary Fig. 4). The amplitude distribution of synthetic SPWs remained close to the training distribution, with a mean difference of 2. 7 detector units and an Earth mover’s distance of 66. 1. Leave-One-Session-Out refitting indicated that the lognormal scale parameter was visibly stable, varying by 9. 5%, whereas the shape parameter fluctuated by as much as 20%. The fitted amplitude distribution captured the pooled data but remained sensitive to session composition.

The generator did not reproduce the full scale of variation in event rates. Resampling the null library alone underestimated between-session variability. Once that variability was added to the prediction interval, the training sessions fell within it except for the SPW rate of Sample 1 (>99th percentile). The held-out Sample 4 matched in SPW rate but fell below the interval in SWR rate and coupling, so the generator reproduces the pooled training regime rather than every session (Fig. 2A). The synthetic SPW inter-event intervals reproduced the right-skewed form and timescale of the empirical data; however, they couldn’t catch the detailed shape of the distribution (*KS* = 0. 173). Therefore, the generator depicted a pooled stationary replay regime rather than a model of animal- or session-specific state fluctuations.

Pharmacological modulation reproduced the direction reported by Kostanšek *et al*. (2023) and estimated the magnitude of antagonist effects at the lowest modeled activation level (OX2R = 0. 3; Table 1). For the OX1R-selective, OX2R-selective, and dual-antagonist models, predicted ripple-incidence multipliers were 0. 53, 0. 38, and 0. 55 (Table 1) relative to baseline, compared with experimental targets of 0. 55, 0. 38, and 0. 51. The OX2R-selective model reproduced reductions in SPW incidence and amplitude together with increased ripple duration under reduced receptor engagement. Across the complete 15-level OX2R grid, 823 of 825 generated SWRs remained within the predefined physiological limits. Mean ripple frequencies remained between 159. 4 and 162. 6 Hz, durations between 49. 6 and 62. 2 ms, cycle counts between 8. 0 and 10. 0, and mean coupling lags between − 4. 6 and + 2. 2 ms. The synthetic libraries therefore preserved physiologically plausible event structure while introducing the OX2R-dependent changes subsequently supplied to the consolidation model.

*Orexin effects dissociate between encoding and consolidation*.

To determine if the non-monotonic DI dependent on OX2R activation was specific to offline consolidation, we separated orexin-mediated effects on EC. Orexin tone varied through encoding, paired with constant consolidation, produced a monotonically increased DI across the tested range (Fig. 3B). Observed mean DI increased from 0. 140 at *ox* of 0. 3 to 0. 161 at 1. 8, with a rank-order relationship between orexin level and agent performance (⍴ = 1. 00). Thus, increasing orexinergic gain throughout encoding produced progressively stronger memory discrimination without an intermediate optimum.

**Figure 3.**
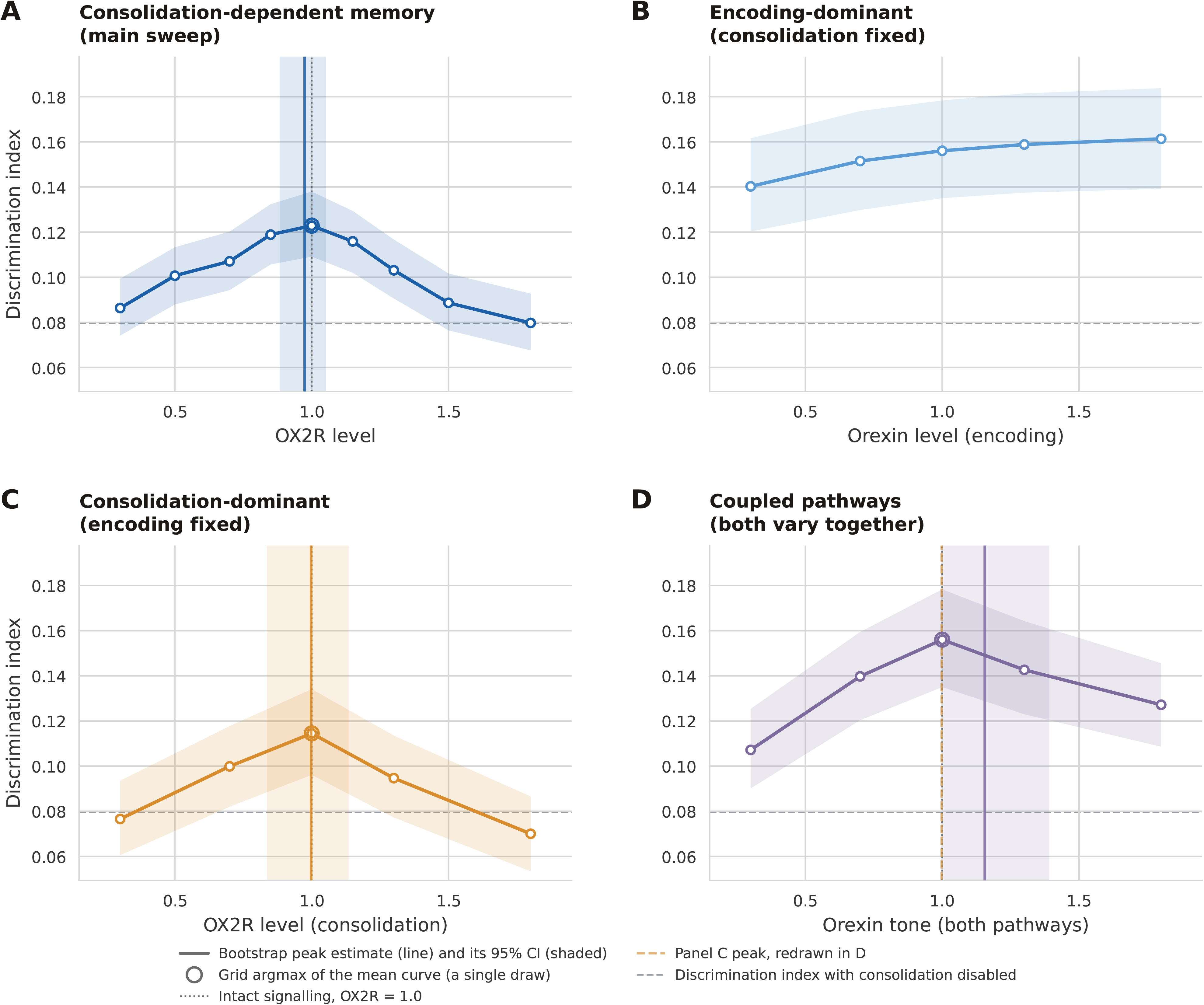
A non-monotonic OX2R–memory relationship appears only when learning depends on consolidation. **(A)** Consolidation-dependent protocol, 9 OX2R levels, 10 encoding presentations. Quadratic coefficient q₂ = −0.0609, P(q₂ ≥ 0) < 10⁻⁴; bootstrap vertex 0.97 [0.88, 1.05], interior in 100% of samples. Both limbs are separately resolved (U-strength +0.0379 [+0.0333, +0.0427]). The optimum sits +0.0432 above the no-consolidation floor (DI = 0.0797); 88% of that gain is lost at OX2R = 0.3, and all of it at OX2R = 1.8, where DI returns to the floor. **(B)** Encoding-dominant arm: consolidation held at baseline, orexin tone scaling encoding gain. The discrimination index rises monotonically (Spearman ρ = +1.000). Here q₂ = −0.0105 reflects saturation rather than a turning point, and the vertex (1.24 [0.30, 1.80]) spans the whole grid and is interior in 0% of samples, so we mark no optimum. **(C)** Consolidation-dominant arm: encoding gain fixed, OX2R acting through replay termination and the induction gate. q₂ = −0.0607, vertex 1.00 [0.84, 1.14], P(interior) = 1.00, ρ = −0.300. **(D)** Both pathways are driven together. q₂ = −0.0602, vertex 1.16 [1.00, 1.39], P(interior) = 1.00, ρ = +0.300; the optimum is displaced +0.158 OX2R units relative to (C). The contrast between (B) and (C) is tested as the OX2R² × arm interaction in a joint fit, DI = b₀ + b₁x + b₂x² + b₃R + b₄Rx + b₅Rx²: b₅ = −0.0502 [−0.0577, −0.0431], P < 10⁻⁴ (paired bootstrap over replicate indices, 10,000 resamples). Points are condition means (n = 50 replicates in A; 25 per level on the 5-level grid in B–D) simulated under common random numbers; shading is the 95% bootstrap CI of the mean, not replicate spread. Vertical band, 95% CI of the bootstrap vertex; open circle, grid argmax; dashed grey line, discrimination index with consolidation disabled. (A) and (C) are the same manipulation on 9- and 5-level grids and are not independent replications.

In contrast, varying effective OX2R activation during consolidation while holding encoding constant produced a curved response (Fig. 3C). DI increased from 0. 077 at OX2R = 0. 3 to 0. 115 at OX2R = 1. 0, before declining to 0. 070 at OX2R = 1. 8. The fitted relationship showed a negative quadratic coefficient (*q*_2_ =− 0. 0607), with a continuous peak at approximately OX2R = 1. 00 and an interior maximum in all bootstrap samples. The main consolidation sweep showed the same pattern, with an estimated optimum of OX2R = 0. 97 ( 95% CI, [0. 88, 1. 05]; Fig. 3A).

When both pathways varied simultaneously, the non-monotonic consolidation profile was retained but shifted toward slightly higher orexin levels (Fig. 3D). The coupled-pathway curve remained negatively curved (*q*_2_ = − 0. 0602) with an estimated optimum at 1. 16 and an interior maximum in all bootstrap samples. By comparison, varying encoding while consolidation was fixed at baseline remained monotonic, whereas varying consolidation with encoding fixed reproduced the intermediate optimum. These results show that the effect of orexin on modeled memory depends on the phase of memory processing: increasing orexinergic drive benefits encoding monotonically, whereas OX2R-dependent modulation of offline replay imposes an intermediate optimum on consolidation (b₅, Fig. 3).

Next, we tested whether the consolidation optimum depended on the supervised delta-rule learning signal used in the primary model. We replaced the delta rule with reward-modulated Hebbian plasticity. The output preserved the non-monotonic memory profile, with negative curvature (*q*_2_ = − 0. 0522) and an estimated optimum at OX2R = 0. 93, closely aligned with the delta-rule optimum at 0. 97 (Supplementary Fig. 2). The estimated difference between the two optima was small (Δ = − 0. 040; 95% CI [− 0. 064, − 0. 023]). Both cases demonstrated maximal transmission of discriminative information to the readout near baseline effective OX2R activation. However, both unsupervised rules (Oja and Sanger) delivered less discriminative information to the readout (1.9-2.2 times chance, against 3.6-5.0 times for the Delta and Hebbian rules). Oja’s rule did not convert it into discrimination (DI near 0.012 throughout); Sanger’s rule has no signed readout. The intermediate optimum therefore did not require an error-correcting teaching signal, although its expression as behavioral discrimination depended on a learning rule capable of associating replayed information with memory outcome (Supplementary Fig. 2).

Together, these results demonstrate that orexinergic modulation has phase-dependent effects on modeled memory performance. Increasing orexinergic drive during encoding produced monotonic improvement, whereas OX2R-dependent modulation of offline replay imposed an intermediate consolidation optimum. The persistence of this optimum under reward-modulated Hebbian learning further indicates that the consolidation-specific non-monotonicity is not an artifact of the supervised delta-rule implementation.

### OX2R-dependent replay dynamics generate a plasticity–truncation trade-off

Varying effective *OX*2*R* activation produced opposing changes in event-level plasticity and replay completeness. At low *OX*2*R* levels, replay sequences were generally untruncated, but the synthetic SWR library supplied fewer events with weaker SPW amplitudes. In the main sweep, the mean number of planned replay events increased from 50 at OX2R = 0. 3 to 107 at OX2R = 1. 0, while the mean drive gate increased from 0.184 to 0.500 and the resulting LTP gate from 0.160 to 0.418 (Fig. 4B). Replay quality didn’t change at low activation scale ( *mean quality gate* = 1. 00 at OX2R = 0. 3∼0. 7). The rise of mean plasticity induction from 0. 087 at OX2R = 0. 3 to 0. 600 at OX2R = 1. 0, and cumulative induction mass from 3. 00 to 23. 73, indicated that the ascending limb was driven by greater effective plasticity rather than by changes in replay completeness.

**Figure 4.**
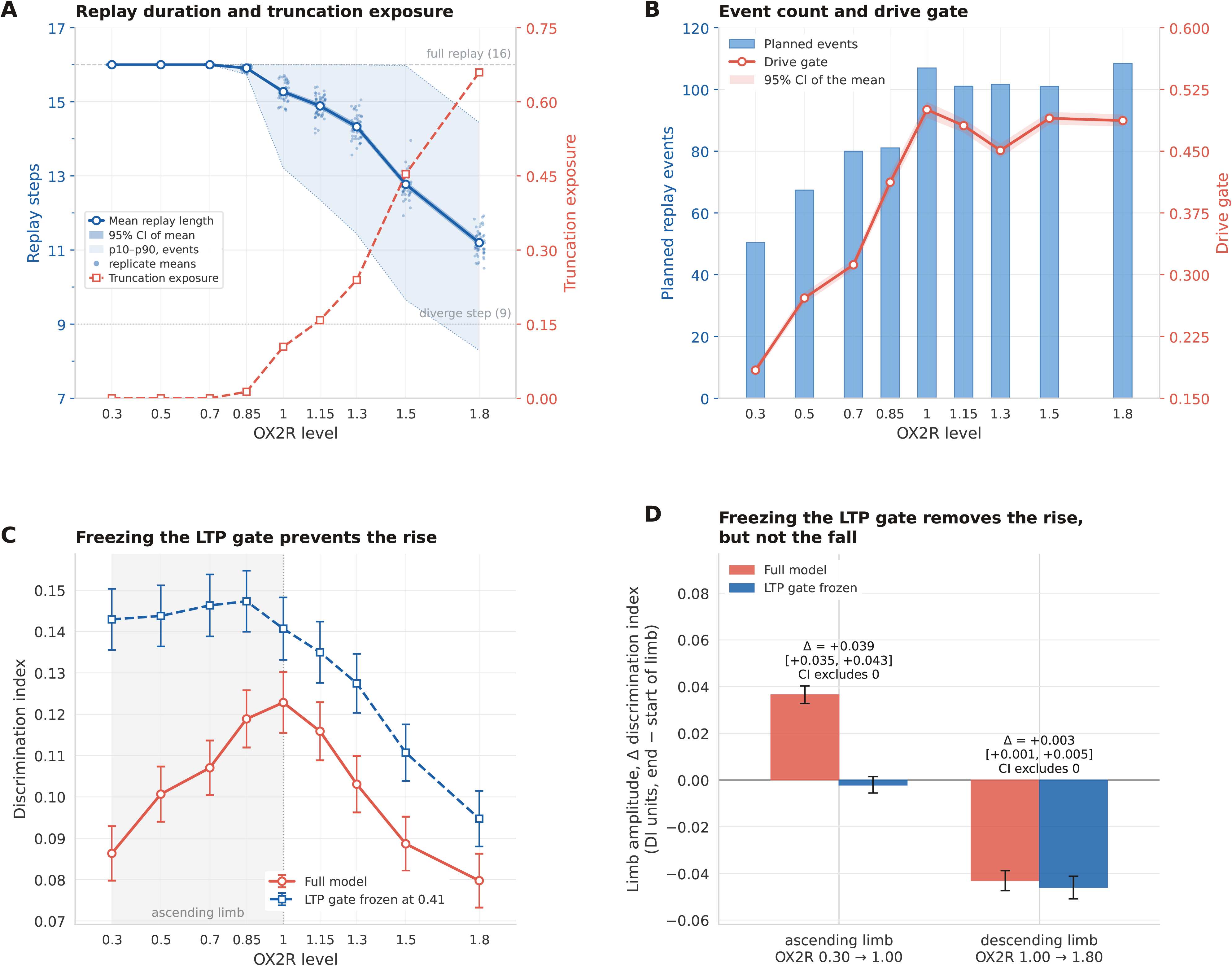
The rise and the fall of the memory curve come from different mechanisms. **(A)** Mean replay length (left axis) and truncation exposure (right axis). Replay runs to the full 16 steps up to OX2R = 0.70 and shortens above it, reaching 11.20 steps [11.10, 11.29] at OX2R = 1.80, while truncation exposure rises from 0.000 to 0.660 across the same range. The pale band is the p10–p90 spread across replay events: at OX2R = 1.80 the tenth percentile is 8.3 steps, below the divergence step (9, dotted line), so part of the event population terminates before target and lure separate and carries no discriminative information. **(B)** Planned replay events (bars, left axis) and the LTP drive gate (right axis). Both rise from OX2R = 0.3 to 1.0 and plateau above it (planned events 50 to 107; drive gate 0.184 to 0.500, 0.487 at 1.8). **(C)** Discrimination index for the full model and with the LTP gate frozen at 0.415. Freezing removes the rise. The frozen arm sits above the full model at every level (mean offset +0.0295), which follows from averaging a thresholded induction function at a fixed gate (mean induction mass 27.22 against 12.00); the panel therefore compares curve shape rather than level, and the limb amplitudes in (D) subtract the offset by construction. **(D)** Limb amplitudes, end minus start of limb, with 95% bootstrap CIs. Ascending limb: +0.0365 in the full model against −0.0022 with the gate frozen, Δ = +0.0387 [+0.0346, +0.0427]. Descending limb: −0.0431 against −0.0460, Δ = +0.0029 [+0.0005, +0.0051]. Freezing the gate abolishes the rise and leaves the fall in place; its residual effect on the descending limb is about one-thirteenth the size of its effect on the ascending limb. The OX2R grid is non-uniform and denser near 1.0 (0.30, 0.50, 0.70, 0.85, 1.00, 1.15, 1.30, 1.50, 1.80), with n = 50 replicates per level simulated under common random numbers. Bands and error bars are 95% bootstrap CIs of the mean unless labeled otherwise. The task has 64 dimensions over 16 replay steps, with 38 shared and divergence at step 9.

During the progressive increase of effective OX2R activation above the baseline, the interaction flipped as replay shortened the available sequence. Thus, mean replay length dropped from 15. 27 steps at OX2R= 1. 0 to 11. 20 steps at OX2R = 1. 8, while truncation exposure went up from 0. 104 to 0. 660 (Fig. 4A). The mean replay-quality gate fell over the same interval. The reduction occurred despite a maintained drive gate at supra-baseline activation; the mean drive gate was 0. 500 at OX2R = 1. 0 and remained 0. 487 at OX2R = 1. 8 (Fig. 4B).

Thus, increasing OX2R beyond baseline did not primarily reduce the amplitude-dependent plasticity drive. Instead, stronger *E*/*I*-dependent termination progressively removed the post-divergence portion of the replay sequence required for discriminative plasticity.

The observed relationship between the plasticity limitation and the replay truncation generated maximal effective plasticity near baseline OX2R activation. Induction mass peaked at OX2R = 1. 0 and then declined sharply to 13. 86 at OX2R = 1. 3, 5. 55 at *OX*2*R* = 1. 5, and 0. 04 at OX2R = 1. 8. The memory performance followed a detected pattern: the DI increased from 0. 086 ± 0. 007 at OX2R = 0. 3 to 0. 123 ± 0. 007 at OX2R = 1. 0, before decreasing to 0. 080 ± 0. 006 at OX2R = 1. 8. The fitted OX2R–performance relationship showed a negative quadratic coefficient (*q*_2_ =− 0. 0609); bootstrap probability of non-negative curvature (< 0. 0001), with a continuous optimum at OX2R = 0. 97 (95% CI, [0. 88–1. 05]) and an interior maximum in all bootstrap samples. The rise from OX2R = 0. 3 to 1. 0 ( Δ *DI* = + 0. 0365) and decline from 1. 0 to 1. 8 (Δ *DI* = − 0. 0431) were comparable in magnitude. Together, these results identify an intermediate regime in which SPW drive is strong enough for event-level induction and replay preserves enough sequence content to support effective consolidation.

### Optimal OX2R activation shifts with task structure and replay demand

We next tested whether the OX2R level that maximized consolidation depended on the depth of replay required to recover discriminative information. Increasing sequence length systematically displaced the optimum toward lower OX2R activation (Fig. 5A). Descriptively, the bootstrapped quadratic vertex declined from OX2R = 1. 51 for an 8-step sequence to 1. 22 for 12 steps and 0. 97 for 16 steps, before falling to 0. 76, 0. 44, and 0. 38 for sequences of 20, 24, and 28 steps, respectively. Several confidence intervals reached the sampled OX2R boundaries of 0. 3 or 1. 8, indicating that those vertex estimates were censored. We therefore treated peak locations as descriptive and tested the shift using a censoring-free interaction model fitted across all OX2R levels and replicates. The OX2R-by-sequence-length interaction was negative (β =− 0. 00718; 95% CI [− 0. 00823, − 0. 00618]; *P*[β < 0] = 1. 000), while the quadratic term remained negative (β_2_ =− 0. 02999; *P*[β_2_ < 0] = 1. 000). The fitted interaction implied a shift of − 0. 0372 OX2R units per additional sequence step (95% CI [− 0. 0406, − 0. 0341]; *P*[*s*ℎ*ift* < 0] = 1. 000). A separate vertex-based analysis showed the same direction descriptively (*slope* =− 0. 0588, 95% CI [− 0. 0815, − 0. 0223]; *P*[*slope* < 0] = 0. 998).

**Figure 5.**
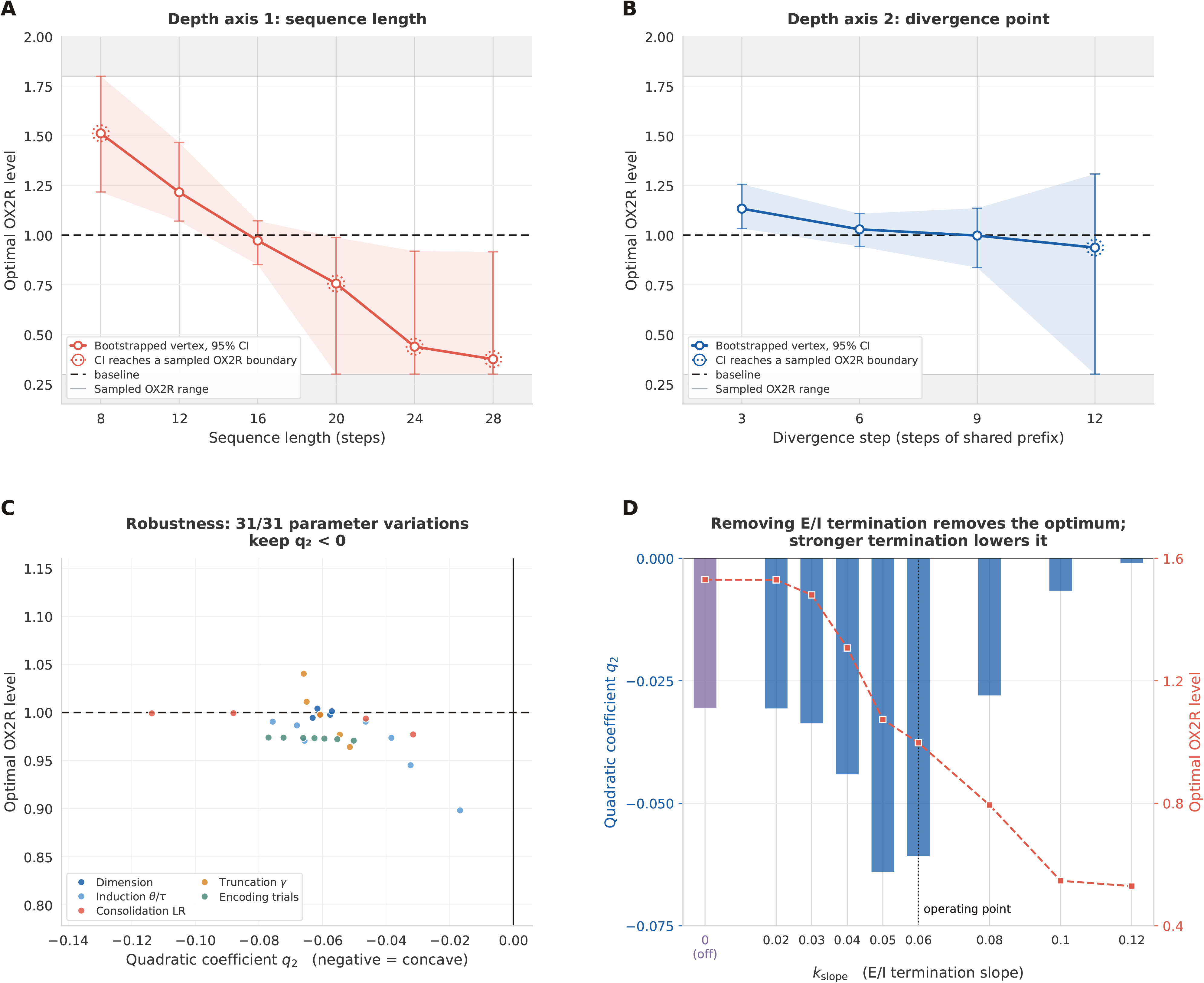
The optimal OX2R level shifts with task demand, and the shift does not depend on vertex location. **(A)** Bootstrap vertex against sequence length (8–28 steps). The estimate moves from OX2R = 1.512 [1.217, 1.800] at length 8 to 0.375 [0.300, 0.916] at length 28, but four of the six intervals reach the sampled floor or ceiling and are censored (dotted ring). The primary statistic is therefore the OX2R × depth interaction, fitted to every replicate at every level: β = −0.00718 [−0.00823, −0.00618], P(β < 0) = 1.000, concavity β₂ = −0.02999, P(β₂ < 0) = 1.000. Implied shift −0.0372 OX2R units per step [−0.0406, −0.0341]. **(B)** The same analysis against divergence step (3–12): vertex 1.133 [1.033, 1.256] at step 3 to 0.937 [0.300, 1.307] at step 12; β = −0.02079 [−0.02269, −0.01890], P(β < 0) = 1.000; β₂ = −0.07252, P(β₂ < 0) = 1.000; implied shift −0.0137 OX2R units per step [−0.0150, −0.0122]. **(C)** Curvature across five parameter axes: coding dimension (5 conditions), induction threshold and time constant (9), consolidation learning rate (5), truncation exponent γ (5), and encoding presentations (7). All 31 conditions retain negative curvature. The criterion in this panel is the sign of q₂ alone, which is weaker than the interior-optimum test used in Figures 3 and 4, since a saturating monotone response also yields a negative fitted q₂ when its quadratic turns over beyond the sampled range. For that reason, we exclude the E/I termination family from this count, and it appears in (D). **(D)** Mechanism sweeps over the E/I termination slope *k_slope_*, which sets how strongly the race shortens replay. Curvature and vertex location both depend on it: q₂ = −0.0305 with the vertex at 1.530 at *k_slope_* = 0, −0.0607 at 0.998 at the operating point *k_slope_* 0.06, and −0.0009 at 0.530 at *k_slope_* =0.12. P(interior) separates the two regimes: 0.529 at *k_slope_* = 0 (no truncation), rising to 1.000 from *k_slope_*= 0.04 upward. With truncation disabled no event is shortened at any OX2R level, the apparent vertex at 1.530 is not resolved against the sampling ceiling, and the descending limb disappears (DI at 1.80 minus DI at 1.00 = −0.0000 [−0.0025, +0.0024], against −0.0431 [−0.0474, −0.0388] at the operating point). Dotted rings mark vertex estimates whose 95% CI reaches the sampled OX2R floor or ceiling (0.30 or 1.80). Interaction coefficients are fitted to all replicates at all levels and never locate a vertex, so they are unaffected by those boundaries. All intervals are 95% paired bootstrap.

To determine whether this effect reflected replay depth itself rather than sequence length specifically, we independently varied the divergence point while keeping total sequence length fixed (Fig. 5B). Moving the divergence point from step 3 to step 12 shifted the estimated optimum from *OX*2*R* = 1. 13 to 0. 94. The OX2R-by-divergence-depth interaction was again negative (β = − 0. 02079; 95% CI [− 0. 02269, − 0. 01890]; *P*[β < 0] = 1. 000), with persistent negative curvature (β_2_ = − 0. 07252; *P*[β_2_ < 0] = 1. 000). This corresponded to an implied shift of (− 0. 0137) *OX*2*R* units for each additional divergence step (95% CI [− 0. 0150, − 0. 0122]); (*P*[*s*ℎ*ift* < 0] = 1. 000). Thus, two independent manipulations that increased the amount of replay required before discriminative information became available both moved the preferred OX2R regime toward lower activation.

The non-monotonic response was robust to broad variation in model parameters (Fig. 5C). Across all 31 conditions, the optimum stayed between OX2R = 0.90 and 1.04, within 0.1 of the baseline estimate: readout dimensionality, induction threshold or slope, consolidation learning rate, truncation exponent, and number of encoding presentations. The *E*/*I* termination slope was analyzed independently as a mechanistic manipulation. These analyses indicate that the existence of an intermediate optimum did not depend strictly on the baseline parameterization.

We therefore examined the *E*/*I* termination slope separately as a mechanistic manipulation rather than as a robustness test (Fig. 5D). The primary model used *k_slope_*, which scaled the rate according to inhibition recorded during an event. Thus, its zeroing terminates truncation entirely, preventing truncation at or before the divergence point. The proportion of bootstrap places the maximum further from the sampled boundaries, falling from 1.000 at the operating point to 0.529 when termination was disabled. The estimated vertex shifted to 1. 530, staying above the calibration point and against the upper end of the sampled range. Negative curvature persisted (*q*₂ = − 0. 0305), predicting that a quadratic fitted to a monotone saturating response turns over beyond the sampled data, so the sign of q₂ does not by itself identify an interior optimum.

The optimum appeared abruptly rather than gradually as termination strength increased. *P*(*interior*) was 0. 540 at *k_slope_* = 0. 02 and 0. 959 at 0. 03, reaching 1. 000 at 0. 04, by which mean replay quality had reduced to 0. 954. Thus, truncation has to exceed a threshold before it can generate an interior optimum. Below it, the model predicts monotone benefit from increasing orexinergic drive. Above the operating point, the optimum shifted progressively downward, to 0.79 at *k_slope_* = 0. 08, 0. 55 at 0. 10, and 0. 53 at 0. 12, remaining interior in all bootstrap draws. The fitted quadratic coefficient weakened toward zero over the same interval, reflecting the growing asymmetry of the response as the peak approached the lower end of the sampled range rather than a loss of non-monotonicity. The two diagnostics thus dissociate in opposite directions at the two ends of the sweep, which is why the interior-optimum criterion rather than the sign of q₂ was used for characterization. Because *k slope* is a free structural parameter not constrained by the pharmacological LFP measurements used to calibrate the synthetic SWR library, this sweep defines the mechanistic dependence of the prediction rather than an externally validated parameter range.

Together, these results show that the optimal OX2R regime is not fixed. As a task requires information to survive further into the replay sequence, high-OX2R conditions become increasingly penalized by premature termination, shifting the consolidation optimum toward lower receptor activation.

### Gated plasticity supports the ascending limb, while replay truncation preserves the descending limb

To separate the contributions of OX2R-dependent plasticity gating and replay truncation to the non-monotonic memory curve, we repeated the consolidation sweep while fixing the event-level LTP gate at its measured baseline value (*g* = 0. 415) and leaving OX2R-dependent replay plans and termination intact (Fig. 4C). Freezing the gate largely eliminated the ascending limb. In the intact model, DI increased by + 0. 0365 from OX2R = 0. 3 to 1. 0 (95% CI [+ 0. 0328, + 0. 0403]), whereas the frozen-gate condition changed by only (− 0. 0022) over the same interval (95% CI [− 0. 0056, + 0. 0014]). The between-condition difference in ascending-limb amplitude was + 0. 0387 (95% CI [+ 0. 0346, + 0. 0427]), indicating that *OX*2*R*-dependent modulation of the plasticity gate was required for the rise in discrimination toward the intermediate optimum.

In contrast, freezing the LTP gate had little effect on the descending limb (Fig. 4D). From OX2R = 1. 0 to 1. 8, DI decreased by − 0. 0431 in the intact model and by − 0. 0460 when the gate was frozen. The difference between conditions was + 0. 0029 (95% CI [0. 0005, 0. 0051]), a magnitude order smaller than the difference in ascending-limb amplitude (+ 0. 0387) and under 7% of the decline itself. Thus, the decline in memory performance at high *OX*2*R* activation persisted when plasticity drive was held constant, consistent with replay truncation rather than declining LTP drive as the principal constraint on the descending limb.

Freezing the plasticity gate also reduced the overall non-monotonic component of the response: the non-monotonic amplitude + 0. 0398 in the intact model to + 0. 0285 in the frozen-gate condition. Importantly, the frozen-gate arm retained substantial induction activity rather than becoming a no-consolidation control, allowing the comparison to isolate modulation of the gate rather than the presence or absence of consolidation itself. As an internal consistency check, discrimination index closely tracked cumulative induction mass across OX2R levels ( ρ = 0. 967, 95% CI [0. 933, 0. 983]), and their estimated optima differed by only 0.03 OX2R units. Together, these manipulations support separable contributions to the two sides of the memory curve: OX2R-dependent gated plasticity supports the increase toward the optimum, whereas OX2R-dependent replay truncation preserves the decline above it.

### The descending limb is not a calibration or numerical artifact

To test whether the cuspid curve is just an artifact of excessive OX2R learning rate, or other numerical instabilities, we have also ascertained that the maximum update remains substantially below the model’s divergence bound, variability does not increase at high OX2R, and performance approaches rather than crosses the no-consolidation floor. (Supplementary Fig. 3). Instability would in any case not predict the dependence of the optimum on sequence length (Fig. 5A), since the stability threshold is set by step size and input norm and is indifferent to where the divergence point falls. Degradation through interference between overlapping patterns is a separate matter and is not what produces the decline here.

## Discussion

The core objective of this study was to determine the intertwined role of orexinergic modulation and OX2R activation during EC, in order to explain the contrasting results in the literature. Previous studies mainly focused on either of the two aspects separately in the context of sleep deprivation (Deadwyler et al. (2007)), or other stress-inducing scenarios (Zhao *et al*. (2014); Staton *et al*. (2018); Han *et al*. (2020)). None of these studies were explicitly designed to analyze both concurrently, nor did they share the same methodologies: different receptors (OX2R vs OX1R), behavioral paradigms, administration times, and the aforementioned physiological conditions. We summarize the findings below and reframe them within our framework.

Sears et al. (2013) showed that OX1R, but not OX2R, blockade during pretraining impairs conditional fear acquisition (during which encoding dominates), but had little effect on posttraining blockade (consolidation). In our model, this is simulated by a global encoding gain mediated by orexin tone, and a specific, OX2R-driven modulation during consolidation. Figures 3B and 3C partially replicate the study scenario, where orexin tone is swept, but OX2R dynamics are not affected, with overall behavioral performance proportional to the tone.

To investigate what makes encoding performance different from consolidation performance, we defined a complexity metric and tested its impact on consolidation (Figure 5A, 5B). We found that the predicted optimal OX2R modulation decreased with complexity: while the same post-training OX2R adjustment falls on different limbs in the two circumstances and modifies performance in distinct directions, a pretraining modification, acting through an encoding gain that does not depend on sequence length, should move both tasks similarly while preserving their relative ordering.

The asymmetry between EC stages accounts for situations where orexinergic stimulation restores impaired performance but doesn’t change it in the normal baseline condition. The phenomenon can be explained by the ascending and descending limbs shown in Figure 3A and C. For example, Orexin-A reversed spatial learning deficits in pentylenetetrazol-kindled rats (Zhao *et al*., 2014). In our model, increasing orexin level during low-plastic encoding corresponds to the ascending limb, as it rescues detection of early replay divergence.

However, due to the E/I processes by OX2R response, further increases in orexinergic stimulation have detrimental effects (descending limb), which explains why during elevated arousal following traumatic stress, exogenous orexin-A impairs rather than improves memory (Han *et al*., 2020). A heightened state is not affecting encoding abilities (as stimuli are no longer being presented, nor are its memories replayed) but rather failure in discriminating between the new stimulus and previous memories. It should be noted, however, that like in Sears *et al*. (2013), this study also considered OX1R, and our model assigned no consolidation-side role to OX1R.

Recent computational and theoretical work suggests that systems consolidation is constrained by the availability and structure of offline replay (Fiete et al., 2010; Chenkov et al., 2017; Singh et al., 2022; Yang et al., 2024). Specifically for us, OX2R blockade extended ripples by 12% (Kostanšek et al., 2023); therefore, endogenous orexin caps event length, and the model extends that interaction above baseline. This estimation cannot be extrapolated to memory since it was made in ex vivo slices, LFP, and without behavioral validation. However, causal manipulation of event duration in vivo supplies the second link. In freely moving rats, closed-loop optogenetic stimulation triggered on spontaneous ripple onset approximately doubled event duration and improved memory during maze learning, while aborting the late portion of ripples impaired memorization (Fernández-Ruiz et al., 2019). The same approach applied during post-learning NREM sleep enhanced hippocampal and prefrontal memory reactivation and rescued recall of an experience that was otherwise too short to consolidate (Robinson et al., 2026).

To model these, we first need to distinguish two separate phenomena: replay completeness, i.e. at which point a replay terminates, and plasticity efficacy, i.e., the magnitude of synaptic modification induced by an effective replay event, which has been discussed at length already. Subsequently, we adopted E/I termination as the model-specific implementation of replay completeness, and found it to be correlated with the cuspid shape of DI performance by OX2R sweep (Figure 5D). The two limbs of the modeled curve occupy the two ineffective combinations. At low OX2R, replay is complete, but drive is weak (mean quality 1.00, drive gate 0.184); at high OX2R, drive is maintained, but replay is curtailed (drive gate 0.487, mean quality 0.340); and the optimum lies where both are adequate. Although no in vivo experiment has yet manipulated orexinergic signaling and evaluated replay content in the same animals, the existing evidence demonstrates that each link has independent empirical support, which corresponds to the model’s predictions, making the direction of the descending limb the conjunction of two separately measured relationships rather than an unconstrained extrapolation.

We further tested whether the descending limb was an artifact of the calibration boundary or fixed replay statistics. The optimum moved when independent model components were perturbed and did not remain tied to the OX2R=1.0 saturation point. Replay truncation also began below this boundary, and increasing OX2R caused performance to approach the no-consolidation floor rather than fall below it. These controls indicate that the descending limb reflects loss of consolidation from replay truncation rather than calibration or numerical instability (Supplementary Fig. 5).

The central contribution of the model is to make replay completeness an explicit determinant of consolidation strength. In contrast to replay-dependent plasticity rules in which replay is treated as a discrete event, our rule scales synaptic modification not only with the OX2R-dependent learning rate but also with the fraction of the represented sequence that survives within each replay event. Models of replay-dependent consolidation typically treat replay as a discrete occurrence suitable for STDP, so that consolidation scales with the number of events and event duration carries no consequence. Thus, the induction dependence on replay length is a requirement for our hypothesis. In its absence, the model would predict monotone benefit from increasing orexinergic drive regardless of the supplied SWR statistics. Therefore, the previously reported behavioral inconsistencies (Deadwyler et al. 2007, Zhao et al. 2014, Han et al. 2020) would remain unresolved. Instead, we add a double gating: plasticity and replay, which also interplay and in turn affect encoding and consolidation.

This coupling produces a trade-off between two effects of increasing OX2R drive. Higher drive increases plasticity during surviving replay, producing the ascending limb, but also increases replay termination and therefore reduces the fraction of the sequence available for induction, producing the descending limb. The importance of the latter effect depends on task demand. Short replay requirements remain largely intact under moderate truncation, whereas longer required sequences lose a greater fraction of their information. Consequently, increasing replay demand shifts the optimum toward lower OX2R drive. In our model, we could dissociate the two limbs beyond simply flattening the cuspid: by making the OX2R gate response not event-driven, the ascending limb was abolished, but the truncation effects, and thus the descending limb, were not (Figure 4D).

The model makes a number of directly testable predictions. The first one is that the effect of post-learning OX2R blockade should depend on how deeply into the replay sequence the task-relevant information occurs. A single antagonist dose during the post-learning rest period should therefore impair consolidation when the relevant information is represented early in replay, and spare or improve it when successful consolidation requires later sequence elements. The model predicts that this effect should be accompanied by changes in ripple duration and replay completeness, or at a higher level by task complexity: contextual fear would represent the ‘easier’ side of the spectrum because the association is available immediately within the represented experience, so consolidation should tolerate truncated replay and benefit from higher OX2R drive, whereas spatial navigation should be negatively impacted as it requires a substantial portion of the trajectory to be replayed to recover the goal location.

Our second prediction is that pre-training orexin tone manipulation should shift performance in the same direction across tasks rather than produce the predicted crossover. Hippocampal recordings during the post-learning rest period would directly test whether the behavioral effect is mediated by reduced replay duration and completeness, thus providing new measurements that no existing study has made in the same animals.

Conversely, we predict that by maxing the OX2R drive decoupled from the SWR drive, only the ascending limb would be abolished. To verify this, an experiment that independently manipulates SWR drive or replay duration while pharmacologically modulating OX2R could test whether the plasticity-dependent benefit of OX2R activation persists when replay truncation is prevented.

There are several limitations to our study. The first kind is about modeling choices. The effects on sharp waves in slices are not receptor-specific, but only OX2R blockade lengthened ripples. Therefore, our model attributes all consolidation-side modulation to OX2R, despite existing evidence of OX1R signaling in the locus coeruleus. None of the reference studies recorded hippocampal SWR statistics during the post-learning rest period. Thus, the mapping can only be tested using the paired behavioral and electrophysiological design described below. Our model is rate-based and abstracts away the spiking and synaptic details of orexin-dependent hippocampal circuit modulation; hence, the OX2R drive parameter represents relative OX2R-dependent modulation rather than a directly measurable quantity such as receptor occupancy, orexin concentration, or receptor activation.

The second limitation concerns the training datasets: the SWR detection and synthetic replay library were derived from a limited number of NREM recording sessions. Despite our best efforts to ground our model in biologically plausible recordings, they do not capture the full variability of hippocampal SWR statistics across animals, sessions, or behavioral states. For modeling purposes, the SWR generator assumes a pooled stationary regime, which captures the central tendency of the empirical data, but is not plausible due to slow fluctuations in network state, inter-animal differences, and context-dependent modulation of excitability. Future extensions should incorporate larger, more heterogeneous recording sets with explicitly non-stationary replay statistics to test the generality of the inferred plasticity–truncation trade-off.

## Data availability

The hippocampal dataset provided by (Grosmark & Buzsáki, 2016) is available to download on the Collaborative Research in Computational Neuroscience (CRCNS) project website (Teeters & Sommer, 2009) under the “hc-11” label. We provide the code used for batch-downloading and performing the experiments at the following address: https://github.com/erolm-a/orexin-swr-consolidation.

Large language models, including Claude Opus (versions 4.6 - 5), were used to assist with code generation, debugging, and refactoring. All AI-assisted code was reviewed by the authors, and the analysis pipeline was executed and validated by the authors. All numerical results, statistical analyses, and figures reported in this study were generated by the resulting computational pipeline and independently inspected by the authors, who take full responsibility for the code and reported results.

## Contributions

P.S. constructed the study, developed the conceptual framework and computational pipeline, designed and performed the analyses and simulations, generated and interpreted the results, prepared the figures, and wrote the initial manuscript; A.G. provided scientific advice and revised the Introduction, Results, and Discussion; E.T. reviewed and revised the computational code and methodology, validated the analysis pipeline and reported results, contributed substantially to interpretation of the findings and development of the Discussion, and established the public code repository. All authors revised and approved the final manuscript.

## Glossary

OX: Orexin peptide OX1R/OX2R: Orexin receptors

Encoding: the initial process of converting a sensory stimulus into a usable pattern

Consolidation: the stabilization of the first trace left by the encoding for long-term storage

EC: encoding and consolidation

SWR: **S**harp **W**aves and **R**ipples, oscillatory patterns produced by synchronized hippocampal neurons, where sharp waves (sudden depolarizations) are quickly followed by ripples (high-frequency oscillations)

EMPA: Selective OX2R antagonist

