## Supplementary for "A Computational Model Captures Orexin-Dependent Modulation of Memory Consolidation"

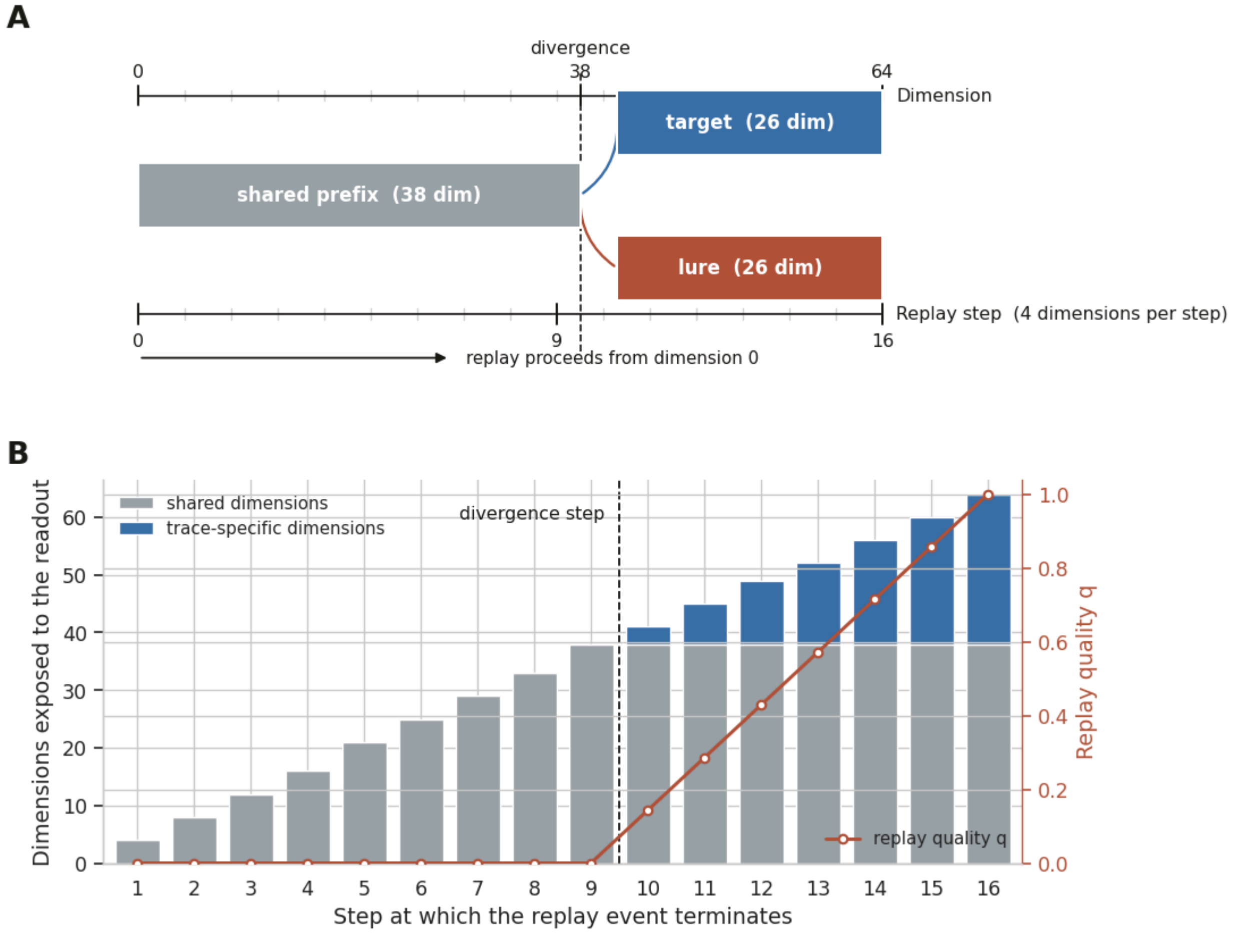


**Supplementary Figure 1. Task structure and replay truncation. (A)** Each trace is a 64-dimensional unit vector whose dimension index stands for position along the remembered sequence. The first 38 dimensions are shared between the two traces; the last 26 are trace-specific. A replay event walks the vector from dimension 0 at four dimensions per step across a nominal 16 steps, so the shared block is exhausted at step 9. **(B)** Dimensions exposed to the readout as a function of the step at which the event terminates, and the resulting replay quality q (right axis), the fraction of trace-specific dimensions revealed. Events ending at or before step 9 expose only shared dimensions and carry no information distinguishing the two traces, however strong the plasticity acting on them; q is zero there and rises to 1 across steps 10 to 16, entering the induction rule directly (Methods, *Consolidation model*). Truncation exponent $\gamma = 1$ throughout the core model.

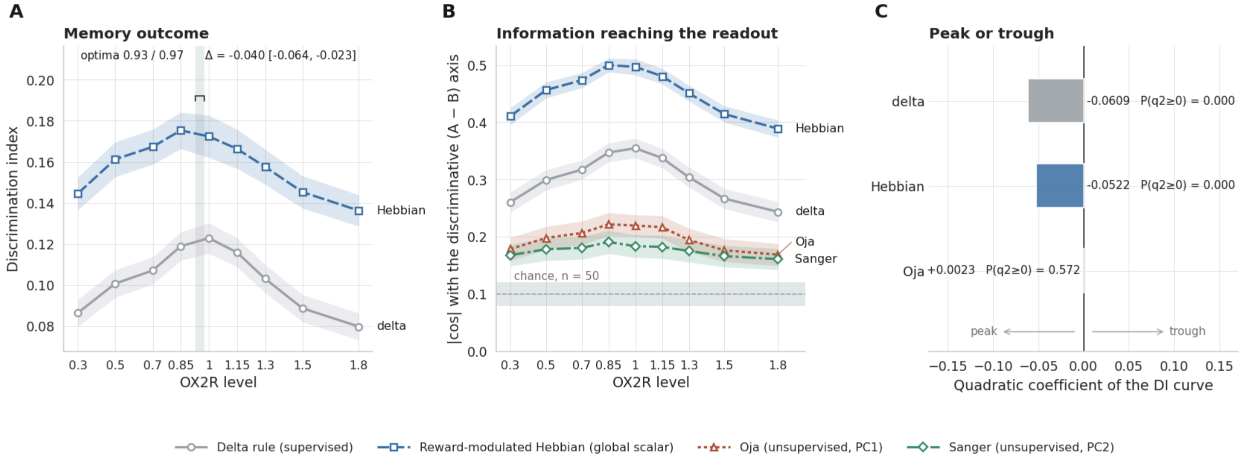


**Supplementary Figure 2. The optimum does not require an error-correcting teaching signal.** Only the readout update differs between rules; sweep, replay plans and seeds are shared. Curves are means ±1 SEM over 50 replicates. **(A)** Discrimination index. Delta and reward-modulated Hebbian rules both peak near baseline (0.97 and 0.93, dotted line; $q_2 = -0.0609$ and $-0.0522$, $P(q_2 \geq 0) < 0.001$), the Hebbian optimum slightly lower ($\Delta = -0.040$, 95% CI [−0.064, −0.023], paired replicate bootstrap). Oja is omitted: a principal component's sign is undetermined and its DI was inconsistently signed across seeds; its coefficient appears in C. **(B)** Alignment between the readout weights and the A−B discriminative axis, |cos θ|. Dashed line and band: chance for a random unit vector in 64 dimensions (0.0997) with its 95% interval (n = 50). All four rules exceed chance and peak near baseline, but the unsupervised rules reach only 2.2 (Oja) and 1.9 (Sanger) times chance against 3.6 and 5.0 for the others. Curvature is reliable for all but Sanger ($q_2 = -0.159$, $-0.156$ and $-0.0754$ for delta, Hebbian and Oja, all $P(q_2 \geq 0) \leq 0.007$; Sanger $-0.0348$, $P = 0.124$). **(C)** Quadratic coefficient of each DI curve; negative is a peak. Oja's is indistinguishable from zero ($+0.0023$, $P(q_2 \geq 0) = 0.572$), with DI near 0.012 throughout, an order of magnitude below delta.

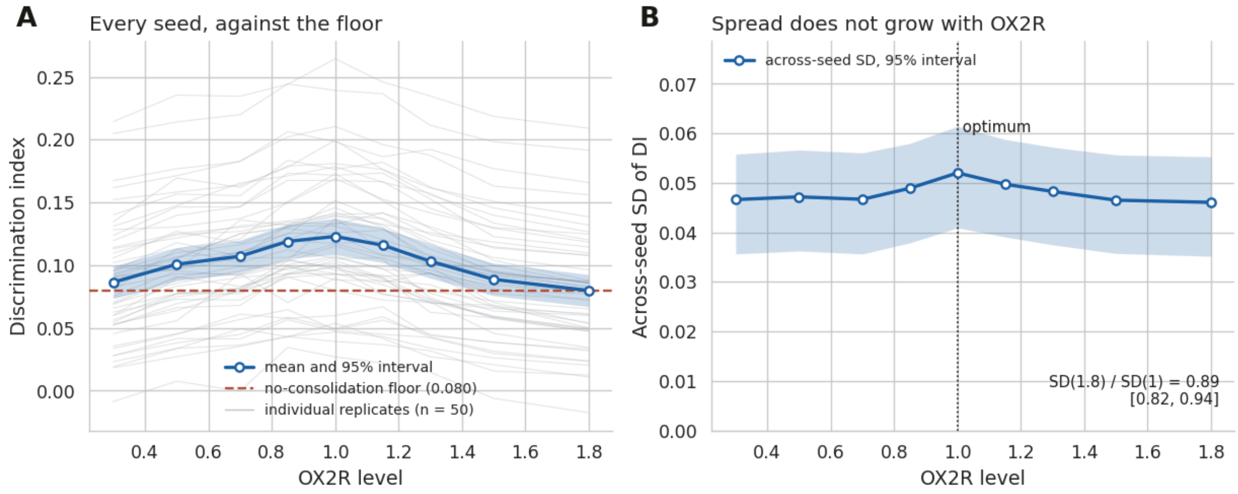


**Supplementary Figure 3. Across-seed spread of the discrimination index does not grow with OX2R.** The main consolidation sweep of Fig. 3A, shown per replicate rather than as a mean: 50 replicates per level under common random numbers, so each replicate contributes the same memory pair, replay plan and trial draws at every OX2R level. **(A)** Discrimination index for every replicate (thin grey), the mean with its 95% interval (blue), and the discrimination index reached when consolidation is disabled (dashed, 0.080). **(B)** Across-seed standard deviation of the discrimination index by OX2R level, with 95% percentile intervals from 10,000 bootstrap resamples of replicates. The standard deviation at OX2R = 1.8 was 0.89 times its value at the optimum (95% CI [0.82, 0.94]; P(ratio > 1) = 0.0002), and the slope of the standard deviation on OX2R was −0.0002 (95% CI [−0.0011, +0.0008]). Step-size instability would show the opposite signature: replicates diverging from one another as OX2R rises, and a mean passing below the no-consolidation floor rather than reaching it.

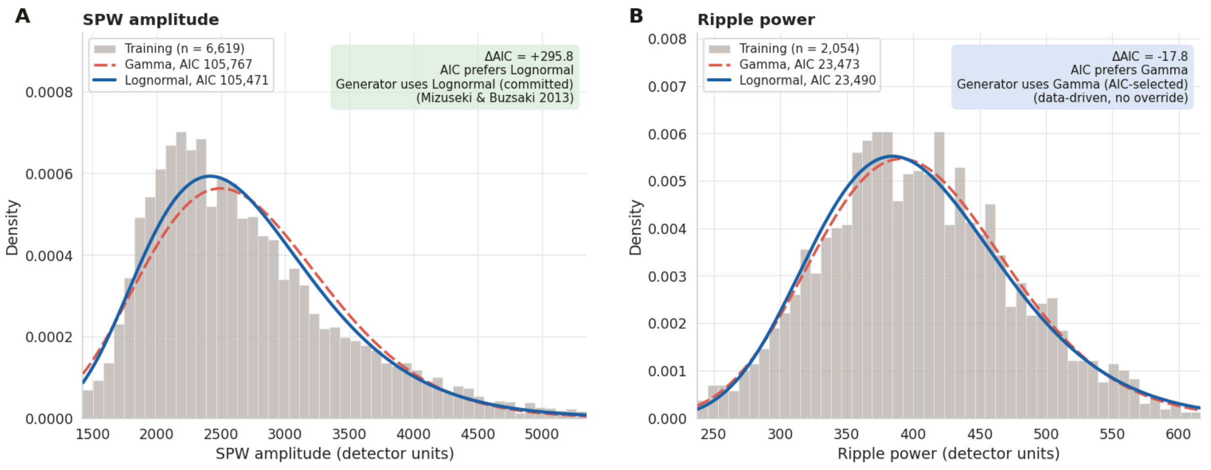


**C**

**Fit comparison**

| | n | Gamma AIC | Gamma KS | Lognormal AIC | Lognormal KS | ΔAIC |
|---|---|---|---|---|---|---|
| SPW amplitude | 6,619 | 105,767.1 | 0.0582 (p = 0.00) | 105,471.3 | 0.0440 (p = 0.00) | +295.8 |
| Ripple power | 2,054 | 23,472.6 | 0.0122 (p = 0.91) | 23,490.4 | 0.0188 (p = 0.46) | -17.8 |

**Supplementary Figure 4. Distribution family for the two event-level amplitude variables.** (**A**, **B**) Gamma and lognormal fits over the histogram of detected events, pooled across the three training sessions; the held-out and excluded sessions are not fitted. (**A**) SPW amplitude (n = 6,619). (**B**) Ripple power (n = 2,054). Both families fix the location at zero and estimate two parameters. (**C**) AIC and one-sample Kolmogorov–Smirnov statistics for each fit. ΔAIC is AIC(gamma) − AIC(lognormal), so positive values favor lognormal and negative values favor gamma. Lognormal fits SPW amplitude better (ΔAIC = +295.8), and the generator samples it as a committed choice, on the reported lognormal statistics of hippocampal magnitudes (Mizuseki & Buzsáki, 2013) as much as on the fit. Gamma fits ripple power marginally better (ΔAIC = −17.8); no literature precedent fixes a family here, so the generator takes whichever family wins the comparison, with no override. For ripple power, neither family is rejected (p = 0.91 for gamma, p = 0.46 for lognormal).

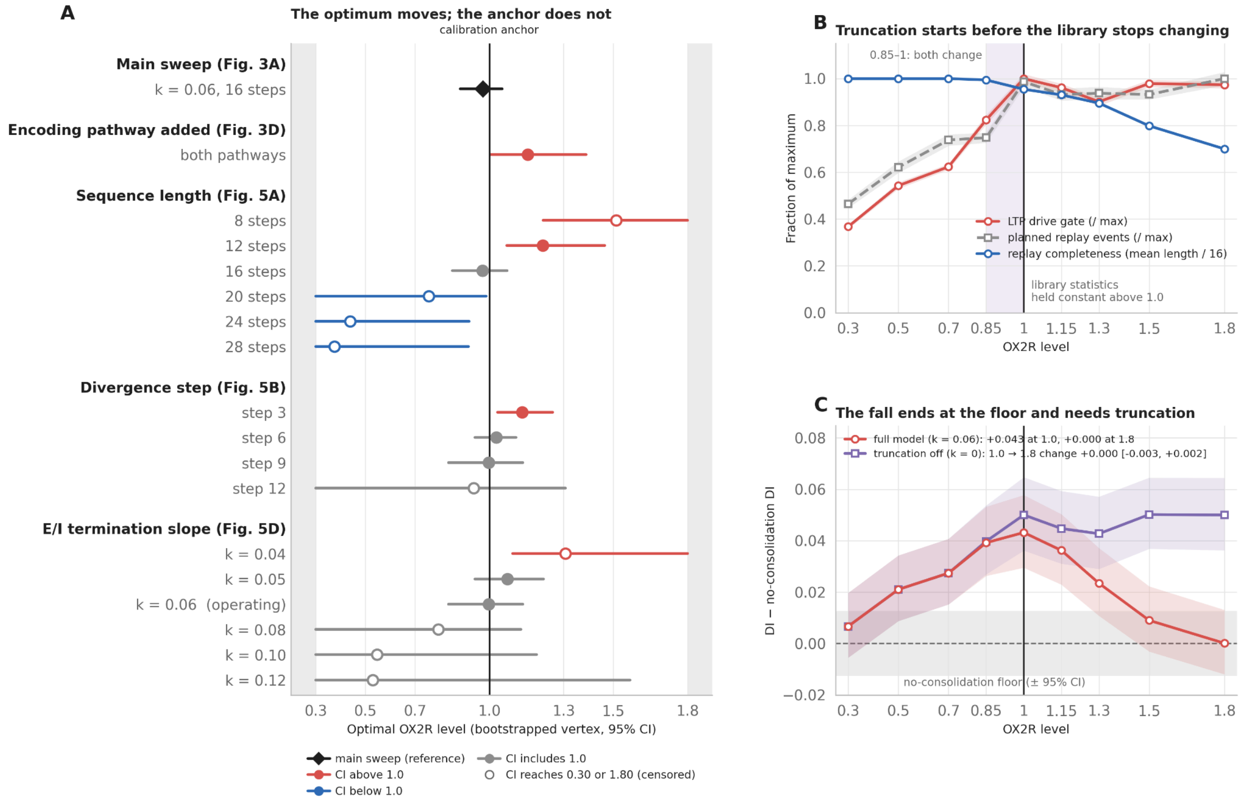


**Supplementary Figure 5. The optimum is not tied to the** OX2R = 1.0 **calibration anchor.** Library statistics were calibrated between the EMPA condition (OX2R = 0.3) and baseline (OX2R = 1.0) and held constant above 1.0. An optimum at 1.0 could therefore reflect where the calibration stops rather than the balance between induction and truncation.

(A) Bootstrapped vertex with 95% CI for each condition, from Figs. 3A, 3D and 5A, B, D. The vertical line marks the calibration anchor, and grey shading lies outside the sampled range (0.30–1.80). Red and blue markers have intervals entirely above or below 1.0, grey markers have intervals that include it, and open markers have intervals that reach a sampled boundary. The main-sweep optimum is 0.97 [0.88, 1.05]. Adding the monotone encoding pathway moved it up to 1.16, a shortened replay sequence (12 steps) to 1.22, and an earlier divergence (step 3) to 1.13 [1.03, 1.26]. Longer sequences moved it down to 0.44 at 24 steps and 0.38 [0.30, 0.92] at 28 steps. Stronger E/I termination lowered it to 0.79, 0.55 and 0.53 ($k_{slope}$ = 0.08, 0.10 and 0.12), although these three intervals include 1.0. An interval censored at 0.30 or 1.80 still excludes 1.0 when its other bound does, since censoring only places the true vertex further from the anchor. The saturation point remained at 1.0 in all conditions, so the optimum could not be displaced within this range if it depended on the saturation point.

(B) Library-side statistics and replay completeness in the main sweep. The LTP drive gate and planned replay events are scaled to their maxima, and replay completeness is mean replay length divided by the 16-step sequence. Above OX2R = 1.0, planned event count stayed between 101 and 108 and the mean drive gate between 0.451 and 0.490. Mean replay length, at the 16-step ceiling up to OX2R = 0.7, had started to decline at 0.85 (15.9 steps, truncation exposure 0.012) and reached 15.27 steps at 1.0. The shaded band marks the range where both change, so truncation and saturation overlap rather than meet at the boundary.

(C) Consolidation benefit: discrimination index minus the discrimination index obtained with consolidation disabled (0.080; grey band, 95% CI of that floor). In the full model ($k_{slope}$ = 0.06), the benefit is +0.043 at OX2R = 1.0 and returns to the floor at 1.8 without passing below it. With E/I termination disabled ($k_{slope}$ = 0), the curve rises to 1.0 and stays flat (DI at 1.8 minus DI at 1.0 = −0.0000 [−0.0025, +0.0024]). Truncation eliminates the advantage of consolidation, so the decline is limited by the consolidation benefit rather than by the termination parameter.

The main sweep and the truncation-off arm use the same 9 OX2R levels and n = 50 replicates under common random numbers, so the two arms share memories, replay plans and trial draws. Vertices in (A) come from the sweeps they cite (n = 25 per level; 5-level grid for divergence step and $k_{slope}$, 9-level grid for sequence length). Bands and error bars are 95% bootstrap CIs over replicates.
